# How the footprint of history shapes the evolution of digital organisms

**DOI:** 10.1101/2021.04.29.442046

**Authors:** Jason N. Bundy, Charles Ofria, Richard E. Lenski

## Abstract

Gould’s thought experiment of “replaying life’s tape” provides a conceptual framework for experiments that quantify the contributions of adaptation, chance, and history to evolutionary outcomes. For example, we can empirically measure how varying the depth of history in one environment influences subsequent evolution in a new environment. Can this “footprint of history”—the genomic legacy of prior adaptation—grow too deep to overcome? Can it constrain adaptation, even with intense selection in the new environment? We investigated these questions using digital organisms. Specifically, we evolved ten populations from one ancestor under identical conditions. We then replayed evolution from three time points in each population’s history (corresponding to shallow, intermediate, and deep history) in two new environments (one similar and one dissimilar to the prior environment). We measured the contributions of adaptation, chance, and history to the among-lineage variation in fitness and genome length in both new environments. In both environments, variation in genome length depended largely on history and chance, not adaptation, indicating weak selection. By contrast, adaptation, chance, and history all contributed to variation in fitness. Crucially, whether the depth of history affected adaptation depended on the environment. When the ancestral and new environments overlapped, history was as important as adaptation to the fitness achieved in the new environment for the populations with the deepest history. However, when the ancestral and novel environments favored different traits, adaptation overwhelmed even deep history. This experimental design for assessing the influence of the depth of history is promising for both biological and digital systems.

## Introduction

In his seminal book *Wonderful Life* [1], Stephen Jay Gould argued for the importance of historical contingencies in the evolution of life on Earth. He proposed a sublime thought experiment of “replaying life’s tape” to ask how similar the living world would be to what we see today if evolution was repeated from, say, the time of the Cambrian explosion. Gould argued that living organisms would likely be very different. However, he concluded, “The bad news is that we can’t possibly perform the experiment.”

But let us presume for a moment that we *could* perform the experiment. Why not up the ante? Why should we rewind the tape to a single point in time, such as the Cambrian explosion some 540 million years ago (mya)? While the *VHS Time Machine* is up and running, why not also dial in some replays from the Paleozoic/Mesozoic boundary (~250 mya) and the late-Cretaceous mass extinction (~65 mya)? By having replays from the dawn of different eras, we could study the influence of varying the time scale over which the signatures of historical contingency become manifest. We could then measure the “footprint of history.” Our own thought experiment thus poses the additional question: “How different would the outcomes be if we rewound the tape of life to different points in history?”

Of course, there was more at stake in Gould’s thought-experiment than biological fantasy. Gould’s idea of “replaying life’s tape” offered a way, however fanciful, to assess the relative importance of the essential components of evolutionary change: selection (adaptation), stochastic variation (chance), and phyletic inheritance (history). So despite its seeming impracticality, an impressive empirical research program has taken up the mantle of Gould’s *gedankenexperiment* [2]. This program includes experimental and comparative studies of the role of contingency and convergence in evolution [3–16]. In particular, the rise of experimental evolution has transformed the idea of “replaying life’s tape” from metaphor to methodology, allowing replay experiments to be conducted in the laboratory using rapidly evolving microbes [2]. While these experiments are not on the grand scale that Gould imagined in *Wonderful Life*, they nonetheless provide a way to estimate the relative contributions of these essential evolutionary components.

Of particular note, Travisano et al. [3] performed a two-phase experiment with bacteria that explicitly partitioned the relative contributions of adaptation, chance, and history to evolutionary change. In the first phase of their experiment, 12 replicate populations of *E. coli* evolved independently for 2,000 generations in a glucose-limited medium [17]. The populations in this phase evolved from the same initial state under identical conditions, thus serving as simultaneous, rather than sequential, replays. The hallmark of Travisano et al.’s experimental design lies in the second phase [3]. A total of 36 clonal isolates (three derived from each replicate in the first phase) evolved for 1,000 generations in a new environment, in which maltose replaced the glucose used in the original environment. Following the second phase, the authors were able to partition the contributions of adaptation (change in the grand mean between the derived Phase II lines and their proximate ancestors), chance (variation among Phase II lines derived from the same ancestor), and history (variation among Phase II lines with different ancestors) to the evolution of both fitness (a trait under strong directional selection) and cell size (a trait not subjected to direct selection) in the new, maltose-limited environment (Fig 1).

**Fig 1.**
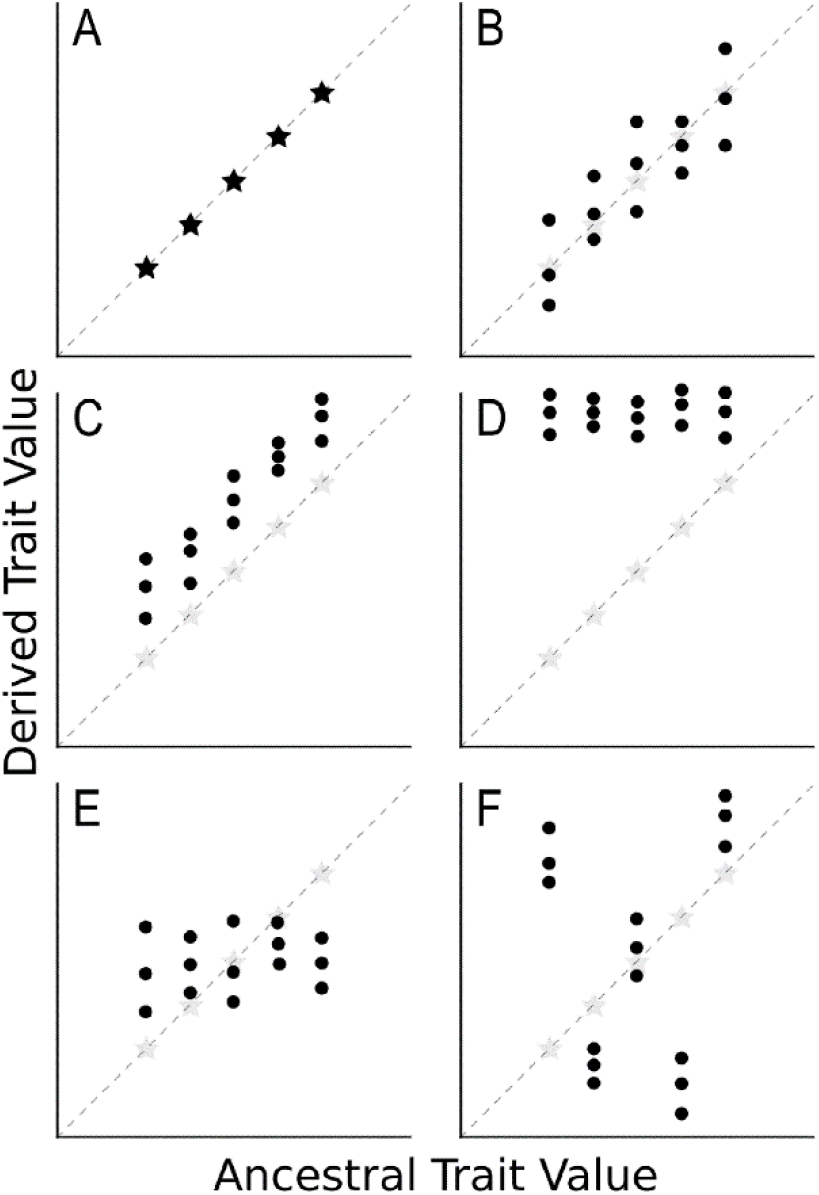
Schematic illustration of the contributions of adaptation, chance, and history. We imagine 5 ancestors arranged along the x-axis according to the mean value of some trait. Each ancestor founds 3 experimental populations that evolve in a new environment, and the mean trait values of the derived populations are shown along the y-axis. **(A)** Initial state: The contribution of history reflects the extent of spread along the x-axis. **(B)** Chance with history sustained: Random factors cause the sets of 3 derived populations that share the same ancestor to diverge from one another, but without any systematic directional tendency. **(C)** Adaptation with history sustained and minimal effect of chance: The trait values of all 15 derived populations have increased (or decreased) systematically relative to their ancestors, and to a similar degree regardless of the ancestral state. **(D)** Adaptation with history erased: The trait values of all 15 derived populations have increased relative to their ancestors, and to a greater extent in those populations founded by ancestors with lower values, such that the effect of history has been eliminated. **(E)** History diminished without directional trend: This pattern might reflect random factors obscuring the effect of history or, alternatively, stabilizing selection on an intermediate trait value. **(F)** History amplified idiosyncratically: Sets of derived populations vary consistently depending on their ancestors, but without any directional trend in trait values or systematic correlation of derived and ancestral values. Modified from Travisano et al. [3].

Experiments of a similar design have now been conducted using dinoflagellates [18,19], an apicomplexan parasite [20], a cyanobacterium [21], the brewer’s yeast *Saccharomyces cerevisiae* [22], and the bacterial pathogen *Acinetobacter baumannii* [23]. These studies have shown that adaptation typically dominates the evolution of fitness and related traits under intense directional selection, whereas chance and history play greater roles for traits subject to little or no selection. As fitness and other strongly selected traits evolve, the effects of the Phase II lineages’ divergent histories—driven by the effects of chance, adaptation, or both in the Phase I environment—were often largely erased [2,3,24–29]. Digital organisms—computer programs that replicate, mutate, compete, and evolve in virtual environments—have also been used to analyze contingency in evolving systems [30–35]. The results of these digital experiments have been consistent with those performed using microorganisms, with adaptation dominating the evolution of traits closely tied to fitness, while chance and history play more prominent roles in the evolution of other traits.

Of particular relevance for our work, Travisano et al. [3] concluded their paper with an intriguing speculation about the possible effect of extending the duration of history in the ancestral environment. Despite having shown that 1,000 generations in maltose (Phase II) largely erased the divergence that took place during 2,000 generations in glucose (Phase I), they suggested: “Over much longer periods, the footprint of history might eventually become too deep to be obscured even by intense selection.” Comparative evidence supports the idea that convergent evolution is less likely when starting from more deeply divergent states [2]. To our knowledge, however, no experiment has systematically quantified the effects of varying the duration of evolution in an ancestral environment—in essence, varying the depth of the footprint of history. Our study addresses this limitation by means of appropriately designed experiments using digital organisms.

### Experimental Design

We performed a set of two-phase replay experiments to test the effects of varying the depth of history in an ancestral environment on subsequent evolution in a new environment. We ran our experiments using the Avida platform for digital evolution [32,36–38]. In Phase I, we started 10 populations from a single proto-ancestral clone that could self-replicate, but which could not perform any of the logic functions that may increase growth rates in their virtual world. These 10 populations evolved independently, but all in the same ancestral (“old”) environment, for 500,000 updates (the basic time unit in Avida), which allowed tens of thousands of generations (mean = 66,074 generations). In this ancestral environment, digital organisms received resources that increased their growth rates if they performed any of 38 distinct two- or three-input logic functions. We isolated the most abundant genotype from each population at three time points, representing three depths of history: 20,000 updates (“shallow”), 100,000 updates (“intermediate”), and 500,000 updates (“deep”). These 30 clones (10 populations × 3 depths of history) were used to seed the Phase II populations.

In Phase II, each of the 30 proximate ancestors identified in Phase I founded 10 new populations in each of 2 new environments, for a grand total of 600 populations (30 clones × 2 environments × 10 replicates). We evolved each of these populations for 100,000 updates, which was equivalent to the intermediate depth in Phase I. One Phase II environment, which we call “Overlapping”, provides the digital organisms with additional resources for performing 76 two- and three-input logic functions, including the 38 functions rewarded in the old environment as well as 38 additional functions. The other Phase II environment, called “Orthogonal”, rewards those 38 additional, novel functions but does not reward the functions that were available in the ancestral environment. We then used the analytical framework of Travisano et al. [3] to calculate the contributions of adaptation, chance, and history to the evolution of average fitness and genome length in each new environment. We are particularly interested in examining the effects of deepening the footprint of history on the relative contributions of those factors, and in testing whether the effects differ between the two new environments (Fig 2).

**Fig 2.**
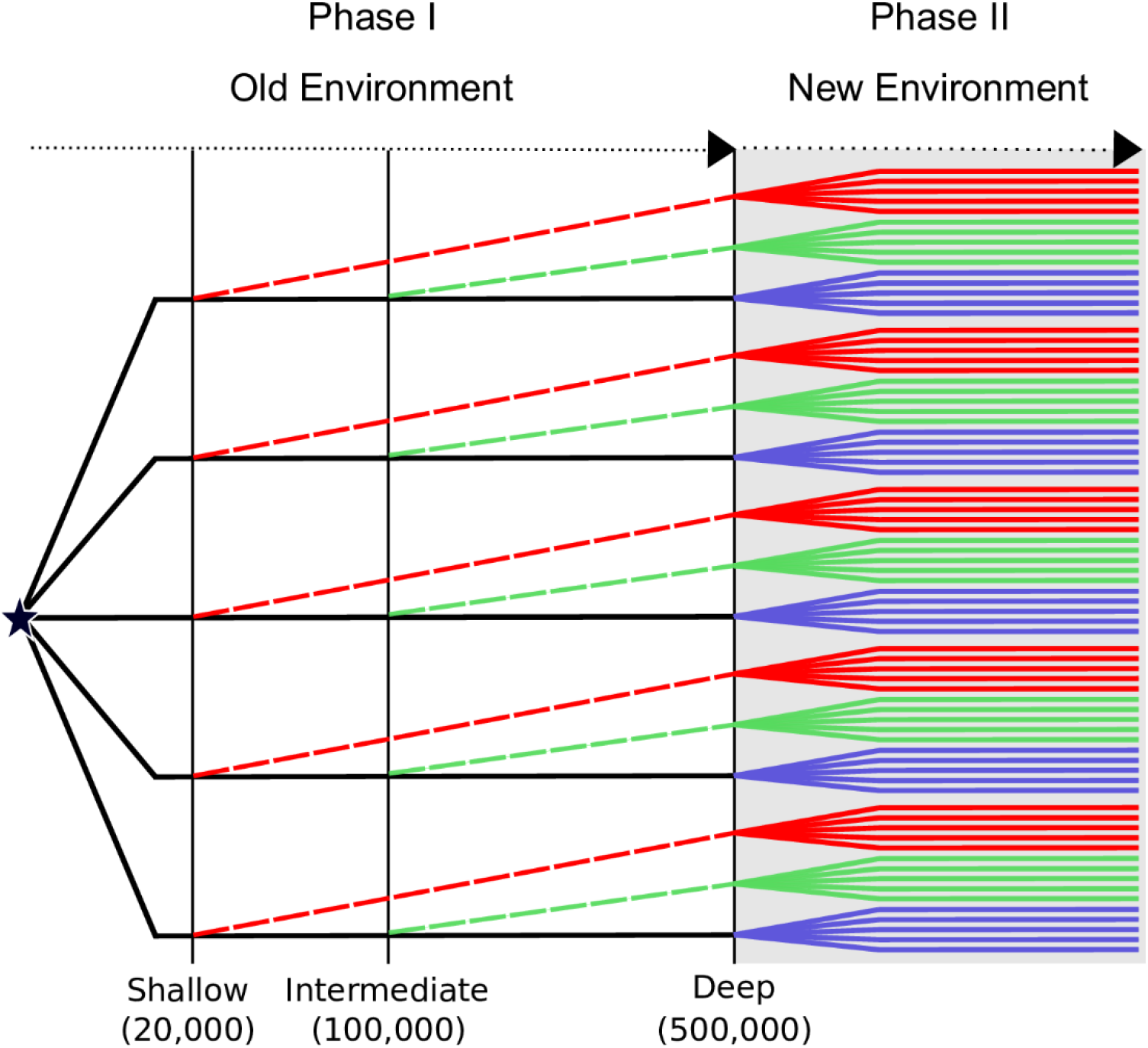
Schematic illustration of the experimental design. In Phase I, ten populations (five depicted) evolved from a single proto-ancestor (star) for 500,000 updates. We sampled the most abundant genotype from each population after 20,000, 100,000, and 500,000 updates, representing shallow, intermediate, and deep history, respectively. In Phase II, each of these 30 proximate ancestors founded ten replicate populations (five depicted) in each of two new environments (one depicted), where they then evolved for 100,000 updates.

In sum, we track 300 evolving populations in each of two environments. Each set of 300 populations has three cohorts (each 100 populations) defined by the depth of their history, with ten founders in each cohort and ten replicate populations per founder. Within each cohort, significant differences in the overall average trait value between replicates (at the end of Phase II) and their direct ancestors (at the outset of Phase II) reflect *adaptation*. For each group of ten replicate populations, evolution proceeds from the same starting point under identical conditions, with only *chance* to explain variation among the replicates. Each lineage’s unique *history* during Phase I accounts for any significant differences among the groups with different founders. Lastly, the *depth of the historical footprint*—reflecting the duration of evolution in the ancestral environment—is responsible for any significant differences among the three cohorts. By employing the analytical methods of Travisano et al. [3], we can estimate the relative contributions of adaptation (systematic changes in mean trait values), chance (variation among replicates), and history (variation among groups with different founders within a cohort). We then compare the contributions among the shallow, intermediate, and deep cohorts to assess the effect of varying the depth of history. We also compare results from the Overlapping and Orthogonal environments to explore whether the impact of that historical footprint depends on the degree of novelty imposed by the new environment.

## Results

### Evolution in Phase I

Starting from the same proto-ancestral genotype, all 10 populations underwent extensive evolution during Phase I. Mean fitness rose rapidly at first, and more slowly as time progressed (Fig 3). The populations all improved relative to their common ancestor, but they reached very different fitness levels (note the log_10_ scale), with the among-population variation increasing over time. The vertical lines show the time points (20,000, 100,000, and 500,000 updates) at which we sampled individual genotypes to start the Phase II populations. We also measured average genome length in each population. Although some variation in genome length emerged over time, there was no consistent directional change: 7 populations finished with slightly longer genomes than the ancestor, 3 ended with somewhat shorter genomes, and the direction of change in an individual population often shifted over time. Moreover, the changes in genome length and the variation among the evolving populations were very small when compared to the fitness trajectories (keeping in mind the different scales for fitness and genome length). Taken together, these observations indicate that genome length was, if not strictly neutral, at least subject to much weaker selection than fitness.

**Fig 3.**
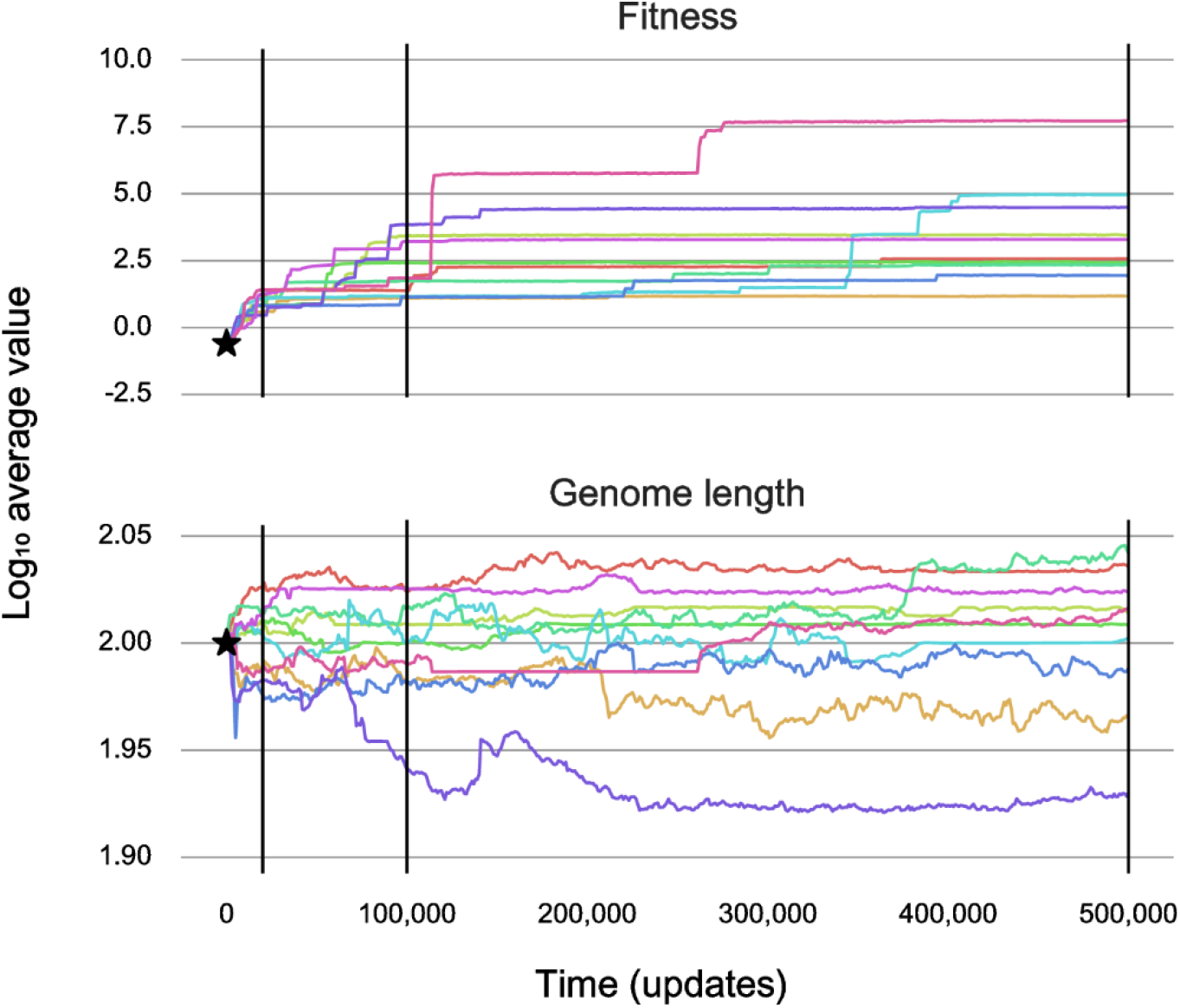
Population averages for fitness and genome length during Phase I. Average fitness (top) and genome length (bottom) measured every 1,000 updates for 10 replicate populations derived from a single proto-ancestor (shown by star), as they evolved in the same Phase I environment for 500,000 updates. We log-transformed both fitness and genome length. Vertical lines at 20,000, 100,000, and 500,000 updates show the times at which we chose the most abundant genotype in each population to start the Phase II replicates.

### Phase II: Evolution in the Overlapping environment after a shallow Phase I history

#### Trajectories for fitness and genome length

Fig 4 (top) shows the mean-fitness trajectories for 100 populations in the Overlapping environment, following a shallow history of only 20,000 updates during Phase I. In this environment, the organisms continued to receive rewards for performing the 38 logic functions rewarded in Phase I. This environment also rewarded the performance of 38 additional functions. We plotted 10 sets of 10 trajectories each, colored according to their Phase I progenitors. Fig 4 (bottom) shows the trajectories for average genome length in the same 100 populations. As in Phase I, fitness increased in all the Phase II populations, whereas genome length hardly changed in most populations and showed little evidence of directionality overall.

**Fig 4.**
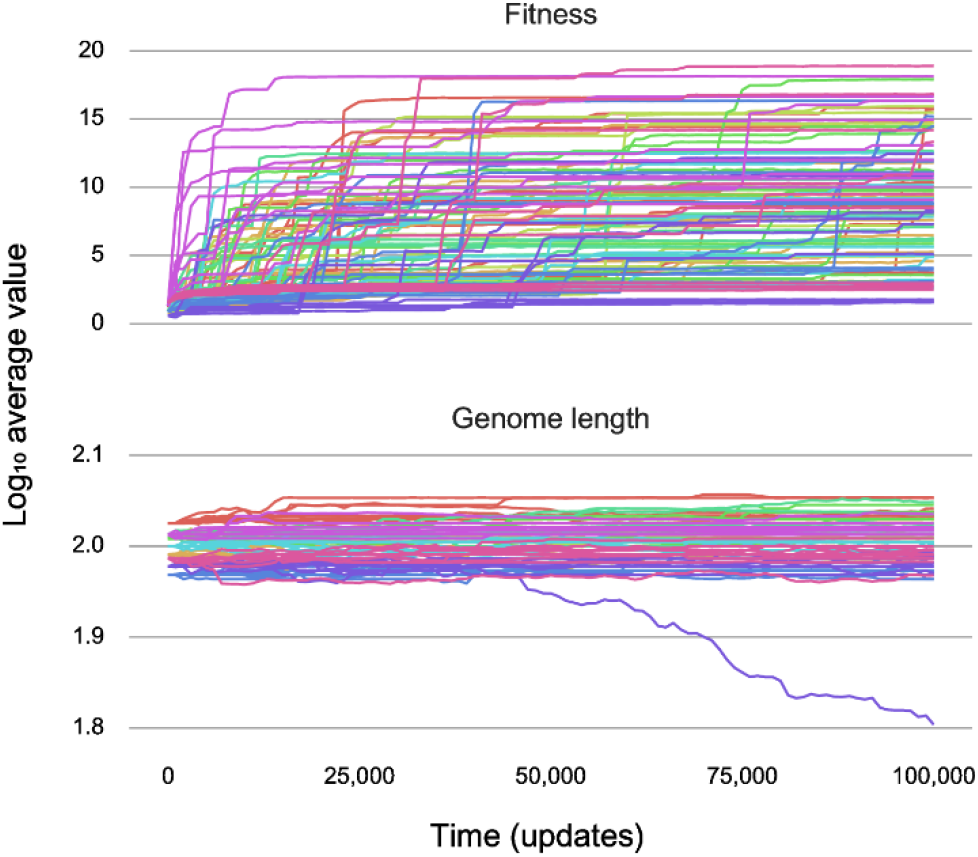
Population averages for fitness and genome length during Phase II in the Overlapping environment for populations with a shallow history. Average fitness (top) and genome length (bottom) measured every 1,000 updates for 10 replicate populations derived from a single founder from each Phase I population, as they evolved for 100,000 updates in the new Phase II environment. These populations have a shallow history, as their founders evolved for only 20,000 updates in the ancestral environment. The new environment rewarded performance of the same 38 logic functions as the old environment along with a suite of 38 additional logic functions.

#### Contributions of adaptation, chance, and shallow history

We can use the data from the endpoints of the evolutionary trajectories to calculate the contributions of adaptation, chance, and history to each phenotype, using the same statistical approach as Travisano et al. [3]. Recall that adaptation represents the overall, directional shift in a trait across populations during Phase II; chance reflects the variation that arose during Phase II among replicates derived from the same Phase I progenitor; and history reflects the variation associated with the different Phase I progenitors. Fig 5 shows these contributions for fitness and genome length after 100,000 updates during Phase II, starting from the 10 progenitors that had evolved for only 20,000 updates in Phase I, thus reflecting a shallow footprint of history. Both panels also show the ancestral variation in those traits among the 10 different progenitors.

**Fig 5.**
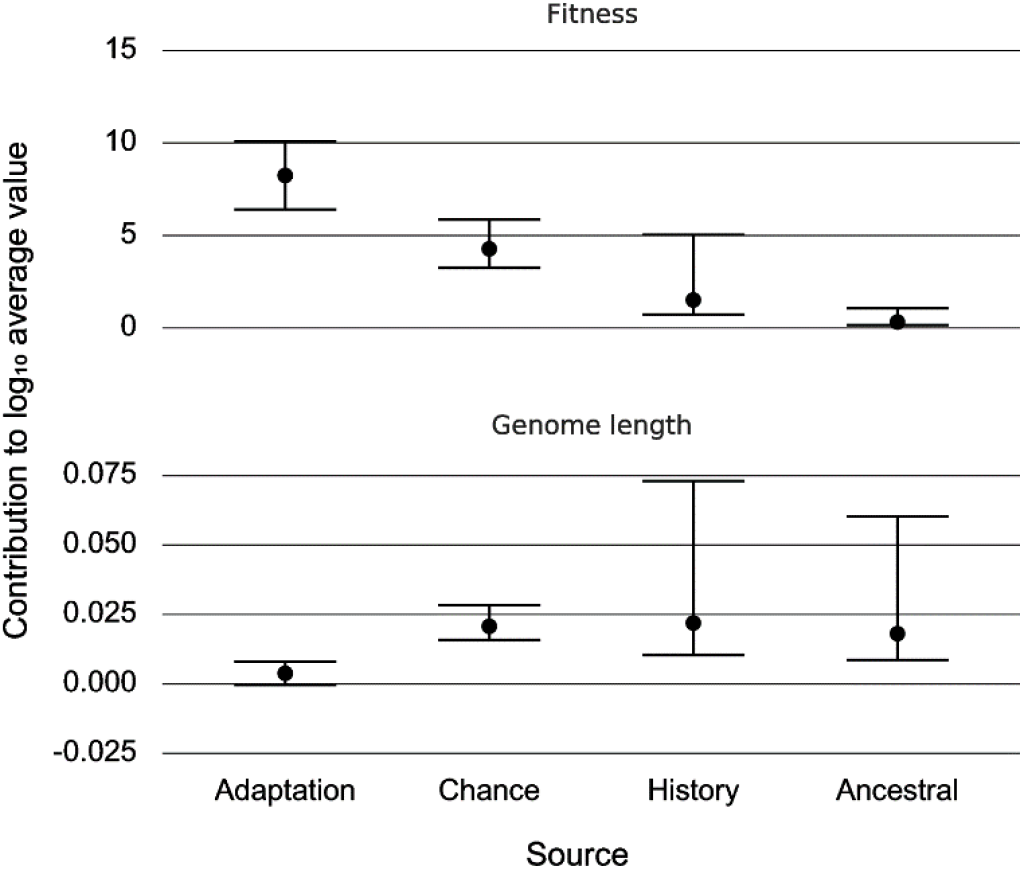
End-of-experiment effects of adaptation, chance, and history in the Overlapping environment with shallow history. Contributions of adaptation, chance, and history to the evolution of fitness (top) and genome length (bottom) based on end-of-experiment average values for populations with a shallow history in Phase I that evolved in the Overlapping environment during Phase II. Error bars show 95% confidence limits for the associated contributions, along with the ancestral variation at the start of Phase II.

In the case of fitness (Fig 5 top), we see a large contribution of adaptation, which reflects the systematic improvement in all 100 Phase II populations. We also see significant, but smaller, contributions that reflect the effects of chance and history. All three contributions, individually and collectively, are greater than the ancestral variation that existed at the start of Phase II. This ancestral variation is the initial Phase II historical variation, which arose during Phase I. For genome length (Fig 5 bottom), by contrast, we see no significant contribution of adaptation (the confidence interval includes 0). However, both chance and history contributed significantly to the final genome lengths, and the variation due to history is similar in magnitude to the ancestral variation in that trait.

We performed these analyses using the values at the end of Phase II, but we also measured the trajectories for all three contributors over time, as shown in Fig 6. We see the rising impact of adaptation on fitness over time, as well as the more slowly rising and meandering contributions of chance and history (Fig 6 top). In the case of genome length, we see the growing impact of chance events, a near-constant effect of history, and a much smaller contribution of adaptation (Fig 6 bottom).

**Fig 6.**
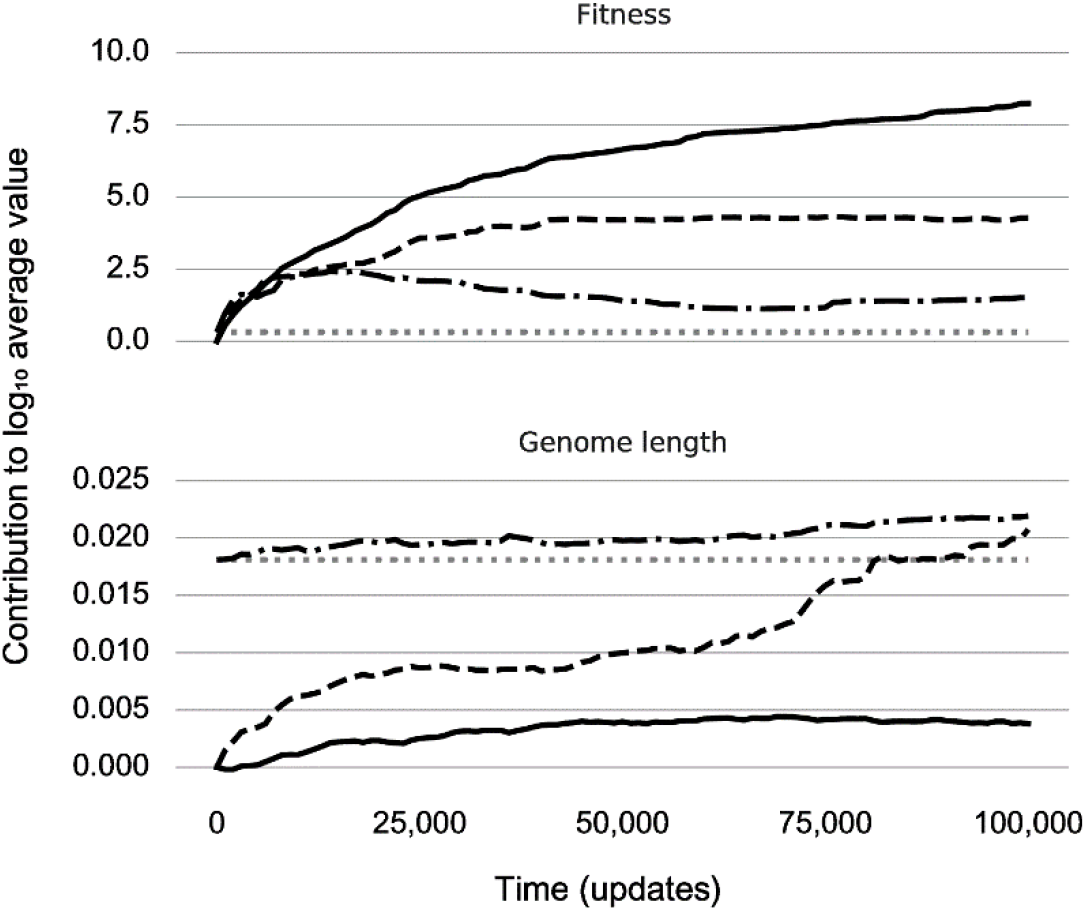
Trajectories of adaptation, chance, and history in the Overlapping environment with shallow history. Contributions of adaptation (solid line), chance (dashed line), and history (dash-dotted line) to the evolution of fitness (top) and genome length (bottom) measured every 1,000 updates during Phase II. The plot includes the ancestral variance (dotted line) for comparison.

### Phase II: Evolution in the Orthogonal environment after a shallow Phase I history

We now turn to the results of evolution in the Orthogonal environment, using the same 10 progenitors that evolved for only 20,000 generations in Phase I. From the perspective of the digital organisms, the Orthogonal environment is more novel than the Overlapping environment. The Overlapping environment continued to reward organisms for performing the same 38 logic functions that the ancestral environment rewarded during Phase I. It also rewarded 38 additional functions exclusive to Phase II. By contrast, the Orthogonal environment rewarded *only* these new functions, and not those rewarded in the ancestral environment. In both new environments, some genetic instructions involved with performing the previously evolved functions might serve as building blocks that could be repurposed to perform the newly rewarded functions. Having led the reader carefully through the data and inferences for the Overlapping environment, we relegate most data and analyses for the Orthogonal environment to the Supporting Information.

#### Trajectories for fitness and genome length

The fitness trajectories in the Orthogonal environment (S1 Fig top) are similar in form to those seen in the Overlapping environment (Fig 4 top). However, the trajectories in the Orthogonal environment started from lower values, and typically reach lower final values, than in the Overlapping environment. The differences between the fitness and genome length trajectories according to environment reflect the continuity of selection. Adaptations acquired during Phase I carried over to the Overlapping environment. However, fitness collapsed in the transition to the Orthogonal environment because the ancestors from Phase I generally lacked the ability to perform functions rewarded in this more distinct, new environment. The trajectories for average genome length in the Orthogonal environment (S1 Fig bottom) were also similar to the corresponding trajectories from the Overlapping environment (Fig 4 bottom). In both cases, most trajectories showed little net directional change, indicative of a trait that is nearly neutral.

#### Contributions of adaptation, chance, and shallow history

As before, we can use the final trait values to calculate the contributions of adaptation, chance, and history to mean fitness and genome length (S2 Fig), as well as the temporal trajectories for these contributions (S3 Fig). For genome length, the contributions of adaptation, chance, and history were nearly indistinguishable in the two environments. In both environments, the lingering effects of history hardly changed over time, while the effects of chance crept upward, slightly eclipsing the effect of history in the Orthogonal environment. Also, adaptation had no significant effect on genome length in either environment. These trajectories indicate a trait that experiences little or no directional selection. The contributions of adaptation, chance, and history to fitness were also similar in the two environments, and their rank orders are the same. However, all the values are substantially larger in the Overlapping environment. Again, this difference in scaling reflects the fact that additional logic functions continued to be rewarded in the Overlapping environment, leading to higher trajectories there (Fig 4 top) than in the Orthogonal environment (S1 Fig top).

Summarizing to this point, we see a clear difference between the contributions of adaptation, chance, and history to the two traits that we measured. For fitness itself, adaptation had the largest effect, followed by chance producing substantial divergence even among the replicate populations founded from the same proximate ancestor. The lingering effect of history, reflecting the different proximate ancestors, was also significant but smaller than the other two effects. For genome length, by contrast, the effects of chance and history were similar in magnitude and both significant, whereas the effect of adaptation, if any, was small and not significant. Also, it would seem thus far that the environment hardly matters for these patterns. We now turn our attention to whether and how these results depend on the depth of the footprint of history.

### Effects of deepening the footprint of Phase I history on evolution during Phase II

The results above are for Phase II populations founded by progenitors that had diverged from the common proto-ancestor for only 20,000 updates. In other words, the footprint of history during Phase I was shallow relative to the duration of Phase II, which was 100,000 updates. Here we examine the effects of history by also using genotypes sampled after 100,000 and 500,000 updates of Phase I to start populations, which then evolved for 100,000 updates in Phase II. In comparison to the shallow footprint of history generated during 20,000 updates, we refer to the footprints that emerged after 100,000 and 500,000 updates as intermediate and deep, respectively.

We provide the underlying data in the Supporting Information, as follows. S4 Fig shows the trajectories for mean fitness in the Overlapping and Orthogonal environments, in each case using 10 founders sampled from Phase I after 100,000 updates (Fig 3). Fig S5 shows the corresponding trajectories for genome length in the two environments. The data in S4 Fig and S5 Fig thus reflect the intermediate footprint of history. S6 Fig shows fitness trajectories in the Overlapping and Orthogonal environments, using 10 founders sampled from Phase I after 500,000 updates; and S7 Fig shows trajectories for genome length in these environments. The data in S6 Fig and S7 Fig reflect the deep footprint of history.

In the two sections below, we examine the contributions of adaptation, chance, and history to genome length and fitness, respectively, comparing the contributions as a function of the depth of history during Phase I and the selective environment in Phase II. We invert the order of presentation and begin with genome length because the patterns and interpretation are simpler than they are for fitness.

#### Effect of deepening the footprint of history on genome length

Fig 7 (top) shows the contributions of adaptation, chance, and history to final genome length as a function of the depth of history during Phase I for the populations that evolved in the Overlapping environment. On the right, one sees the effect of history on the ancestors used to found the Phase II populations; this effect became progressively larger as the depth of history increased from 20,000 to 100,000 to 500,000 updates. To the left of the ancestral estimates, one sees that these historical effects were maintained in the evolving lineages during Phase II. Chance shows the opposite trend, with its contribution declining as the depth of history increased. Adaptation—as indicated by consistent directional changes in trait values—contributed little to genome length in comparison to either history or chance. However, the insignificant contribution of adaptation to genome length in the Overlapping environment when the depth of history was shallow (20,000 updates) became significant when history was deeper (100,000 and 500,000 updates), although the effect remains small. S8 Fig (top) shows the temporal trends in the contributions of adaptation, chance, and history to genome length during Phase II for all three depths of history.

**Fig 7.**
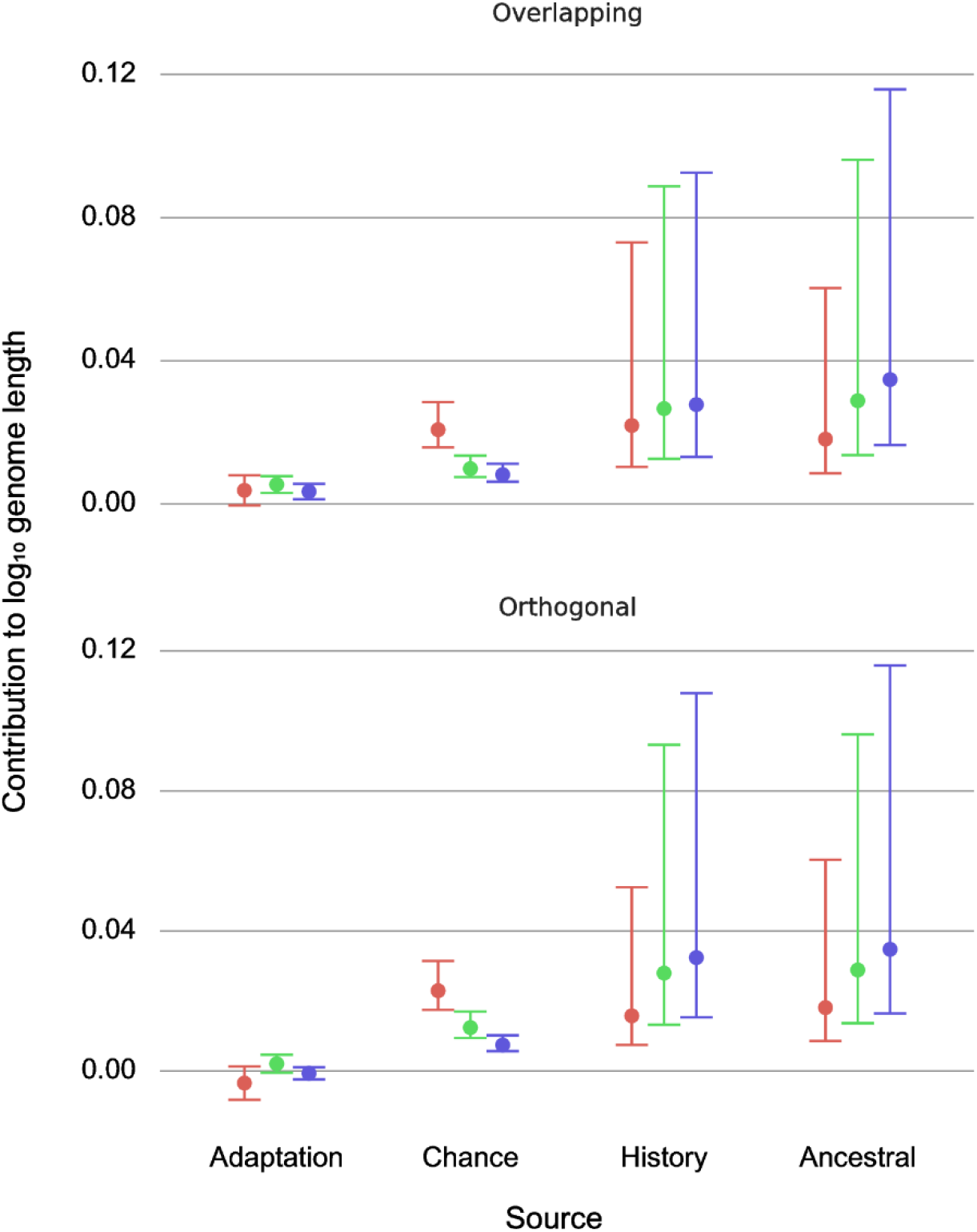
Impact of the footprint of history on the contributions of adaptation, chance, and history to genome length is similar in the two Phase II environments. Contributions of adaptation, chance, and history to genome length for populations with shallow (red), intermediate (green), and deep (blue) Phase I histories. Contributions are based on end-of-experiment averages for populations that evolved in the Overlapping (top) and Orthogonal (bottom) Phase II environments. Error bars show 95% confidence limits. The plot shows ancestral variation at the start of Phase II for comparison.

The end-of-experiment estimates and their trajectories are similar when we look at the effects of the depth of history on genome length in the Orthogonal environment (Fig 7 bottom, S8 Fig bottom). The ancestral divergence increases with the depth of history, as do the persistent contributions of history, while the effect of chance declines as the footprint of history gets deeper. The only qualitative difference in the Orthogonal environment, when compared to the Overlapping regime, is that there was no significant adaptation associated with the deeper histories. Given the lack of continuity in selection for specific logic functions, even the weak selection on genome length that might have occurred in the Overlapping environment was evidently inconsequential in the Orthogonal environment.

#### Effect of deepening the footprint of history on fitness

The results for fitness are more complex, with populations in the two environments showing different effects of the depth of history. Fig 8 shows the contributions of adaptation, chance, and history to final mean fitness for the three depths of history in the Overlapping (top) and Orthogonal (bottom) treatments. In the Overlapping environment, adaptation swamped the effect of history when the footprint of history was shallow. As the depth of history increased from 20,000 to 100,000 and 500,000 updates, the contribution of adaptation declined, while the contribution of history to fitness increased. In fact, for populations with the deepest ancestry, history’s contribution was as great as or slightly greater than that of adaptation, whereas for populations with the shallowest history the contribution of adaptation was several times that of history.

**Fig 8.**
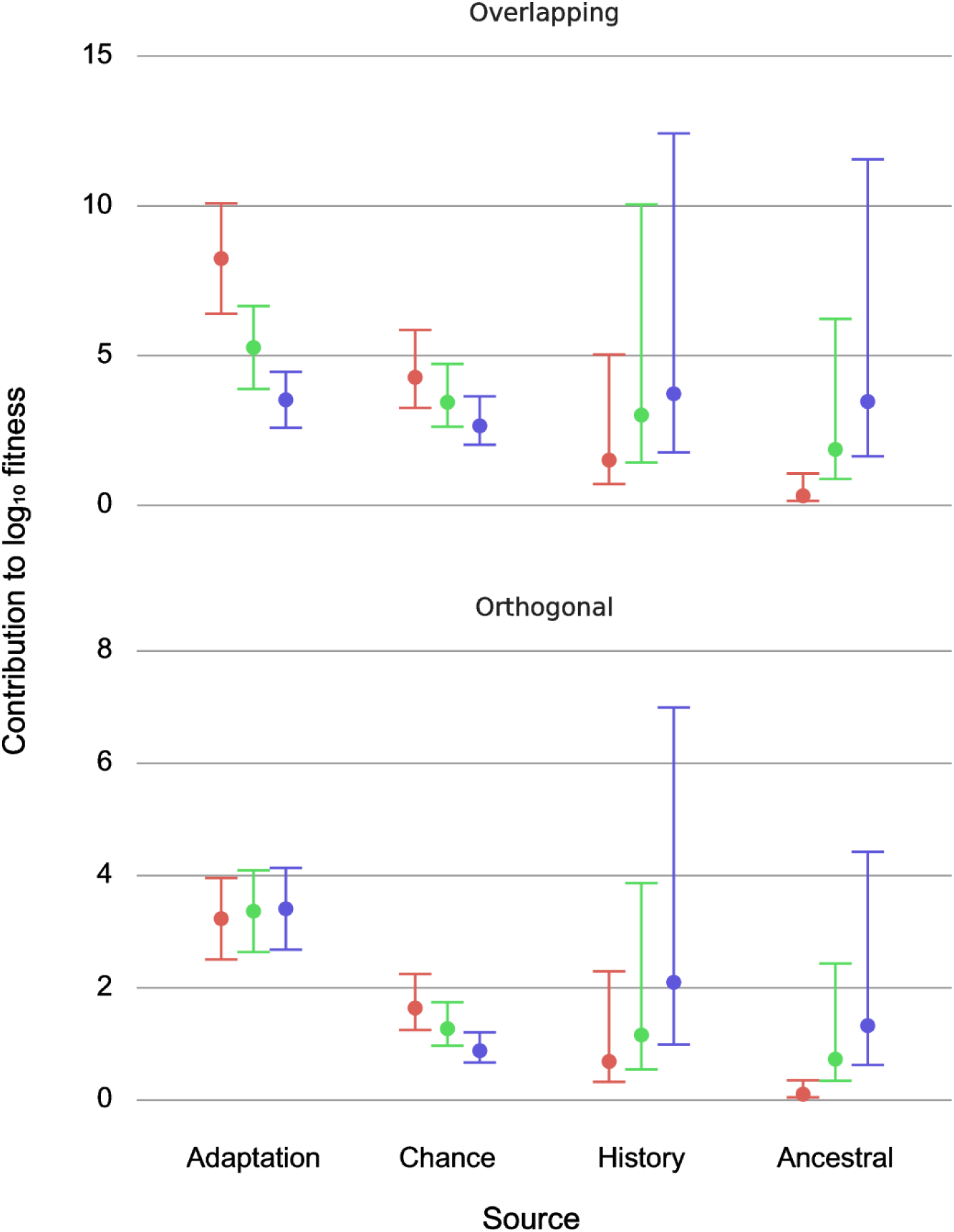
Impact of the footprint of history on the contributions of adaptation, chance, and history to fitness differs between the two Phase II environments. Contributions of adaptation, chance, and history to mean fitness for populations with shallow (red), intermediate (green), and deep (blue) Phase I histories. Contributions are based on end-of-experiment values for populations that evolved in the Overlapping (top) and Orthogonal (bottom) Phase II environments. Error bars show 95% confidence limits. The plot shows ancestral variation at the start of Phase II for comparison.

By contrast, in the Orthogonal environment, where the logic functions rewarded in Phase I were no longer rewarded during Phase II, the contribution of adaptation was essentially independent of the depth of history. The contribution of history to fitness nonetheless increased with its depth, as it did in the Overlapping environment. However, even when the Phase I history was its deepest at 500,000 updates, and thus 5-fold longer than Phase II, the contribution of adaptation to fitness in the new environment remained greater than that of history.

Fig 9 shows the temporal trajectories of the estimated contributions of adaptation, chance, and history during Phase II in the Overlapping and Orthogonal environments. These trajectories further illustrate the impact of the different environments on the relative impact of adaptation and history. Notice that in the Orthogonal environment, the contribution of adaptation surpassed that of even the deepest history within a few thousand updates, whereas it had not done so even after 100,000 updates in the Overlapping environment.

**Fig 9.**
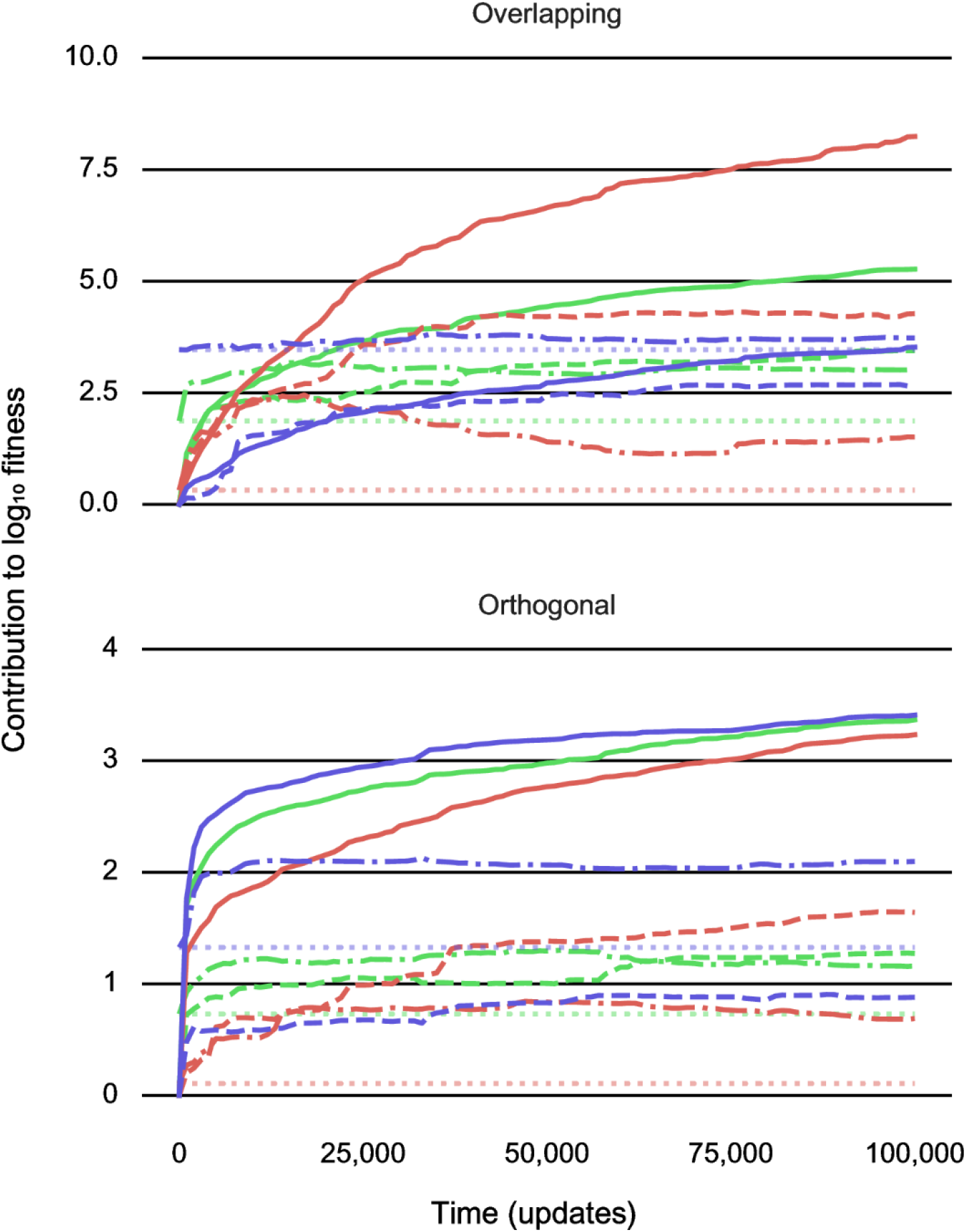
Trajectories for the contributions of adaptation, chance, and history to fitness in the new Phase II environment. Contributions of adaptation (solid lines), chance (dashed lines), and history (dash-dotted lines) to the evolution of fitness in the Overlapping (top) and Orthogonal (bottom) environments measured every 1,000 updates during Phase II for populations with shallow (red), intermediate (green), and deep (blue) histories from Phase I. The plot shows ancestral variances (dotted lines) for comparison.

#### Confidence intervals and confident inferences

The results above are based on a total of 600 Phase II populations (2 environments × 3 depths of history × 100 replicates) and 2 focal traits (fitness and genome length). We did not emphasize statistical details when making comparisons, as we highlighted differences that are generally well supported by our analyses. With respect to genome size (Fig 7), none of the six 95% confidence intervals for the contribution of adaptation overlap with the corresponding point estimate for either chance or history, nor do any of the confidence intervals for chance and history overlap the corresponding point estimate for adaptation. With respect to mean fitness, we saw an interesting interaction between the depth of history in Phase I and the environment during Phase II in terms of the contribution of adaptation to fitness in the new environment (Fig 8). To formalize that interaction, we calculated the following quantity for each Phase I progenitor and its 60 descendant populations: *Y* = (*A* / *B*) – (*C* / D), where *A* is the mean change in log_10_-fitness in the Overlapping environment after Shallow history, *B* is the mean change in the Overlapping environment after Deep history, *C* is the mean change in the Orthogonal environment after Shallow history, and *D* is the mean change in the Orthogonal environment after Deep history. This difference in ratios thus expresses the environment-dependent change in the effect of the depth of history on the contribution of adaptation to fitness in the new environment. Under the null hypothesis of no interaction between the depth of history during Phase I and the environment in Phase II, the expected value of *Y* is 0. Each difference is independent of the others, and all 10 differences are positive, indicating a consistently greater difference in the contribution of adaptation between populations with shallow and deep histories in the Overlapping environment than in the Orthogonal environment. A non-parametric sign test confirms that this interaction is highly significant (two-tailed *p* = 0.0020).

We also performed a meta-analysis of the overall patterns and their consistency with respect to the effects of the depth of the historical footprint. Figs 7 and 8 combined include 16 cases where we estimated four contributions (adaptation, chance, history, and ancestral) for two traits at three different depths of history. If the effects of the depth of history were statistically meaningless (i.e., if the depth of history’s influence was random), then one would expect the estimated contribution for the intermediate depth to fall between the contributions for the shallow and deep histories only one-third of the time. In fact, the contribution at the intermediate depth was between the other two contributions in 14 of 16 cases. The probability of an outcome as or more extreme than that under the binomial distribution with an expected success rate of 1/3 is extremely low (*p* ≈ 0.00001). Even if we exclude the four ancestral cases (because these measurements are the history estimate at the start rather than the end of Phase II), the contributions associated with the intermediate depth fell between the two other contributions in 10 of 12 cases, which remains highly significant (*p* ≈ 0.00054). Only the effect of adaptation on genome length lacks a clear trend with respect to the depth of history, consistent with that trait having been selectively neutral or nearly so. These otherwise consistent outcomes reinforce the striking interaction between the depth of history and the reward structure of the new environment with respect to the contribution of adaptation to fitness (Fig 8).

## Discussion

Gould famously proposed “replaying life’s tape” to gain insight into the role of historical contingency in evolution [1]. Although he viewed this idea as a mere thought-experiment, others developed a quantitative framework for analyzing the contributions of adaptation, chance, and history to phenotypic evolution in experiments with replicated and nested lineages. And while Gould imagined replaying evolution from one moment in time, our experiments were designed to examine how the depth of history affected later evolution. In Phase I, we evolved 10 populations of digital organisms from a common ancestor in a single environment for periods representing shallow, intermediate, and deep histories. In Phase II, these 30 progenitors each founded 10 new populations that evolved in two new environments. Using the statistical framework of Travisano et al. [3], we measured the contributions of adaptation, chance, and history to fitness and genome length. All three factors contributed significantly to the among-population variation in fitness (Fig 8). In the new environment with resources that overlapped those in the old environment, the deep footprint of history contributed as much as adaptation to fitness in the new environment. In the other new environment, which rewarded entirely different functions from the old environment, adaptation overcame even the deep footprint of history. By contrast, history and chance explained most of the variation in genome length, with almost no effect of adaptation (Fig 7). In the future, we hope other researchers will implement our experimental design for analyzing the effects of history’s footprint using biological as well as digital organisms.

Perhaps our most noteworthy observation is that deepening the footprint of history constrained adaptation, but only when the old and new environments were similar. The Phase II Overlapping environment rewarded new functions while continuing to reward those functions that were useful in the ancestral environment. This overlap provided an evolutionary “bridge” that allowed lineages to retain adaptations that arose during Phase I. Deepening history by extending the duration of Phase I provided more time to acquire high-fitness phenotypes in the ancestral environment. These higher—and, importantly, also more variable—initial fitness levels were then sustained in the new Overlapping environment, providing a contribution of history to fitness that increased with the depth of history’s footprint (Fig 8).

By contrast, the Orthogonal environment was largely “blind” to the phenotypic diversity that had been adaptive during Phase I. By chance, some Phase I lineages may have evolved the ability to perform functions that the Orthogonal environment would subsequently reward, even while they were still in the ancestral environment where these tasks were not rewarded. These pre-adaptations could arise as correlated responses because an organism’s ability to perform an unrewarded logic function was connected to its ability to perform a rewarded function. Populations with the deepest ancestry had the greatest opportunity to evolve these cryptic pre-adaptations. Thus, the ancestral variation in fitness at the beginning of Phase II in the Orthogonal environment increased as the footprint of history deepened, just as it did in the Overlapping environment (Fig 8). However, the ancestral variation—even after the deepest Phase I history—was rapidly overwhelmed by selection for new functions in the Orthogonal environment, where the contribution of adaptation was essentially independent of phylogenetic depth (Fig 9).

While adaptation was a potent force driving changes in fitness, that was not the case for genome length, where both chance and history contributed much more (Fig 7). Despite the differential contributions of adaptation, chance, and history to these two traits, it is noteworthy that deepening the footprint of history (by extending the duration of Phase I in the old environment) had consistent ordinal effects on the contributions of adaptation, chance, and history to both traits in each new environment. That is, the intermediate depth almost always yielded contributions between those of shallow and deep history (Figs 7 and 8), showing the importance of the depth of history’s footprint on subsequent evolution. We also observed that adaptation, chance, and history were each capable of being the strongest driver of phenotypic evolution, depending on the specific circumstances of the trait, depth of history, and environment. Experimental approaches to replaying the tape of life thus offer a window into the “blind watchmaker’s” [39] workshop, allowing one to examine how and when various forces contribute to the production of evolved form and function.

As noted by Blount et al. [2], the philosopher of science John Beatty distinguishes between two notions of contingency ascribed to Gould [40]. One is an unpredictability-centered definition of “contingency *per se*” that identifies prior states that are *insufficient* to bring about a given outcome. This version seems to align with the randomness among replicates that we measured as chance. The other is a causal-dependency definition of “contingency upon”, in which certain prior states are *necessary* to bring about a particular outcome. This version aligns more with the lineage effects that depend upon a particular progenitor, which we measured as history. Although these two notions are distinct, Beatty concludes they are compatible. He proposes that contingency might therefore be more fully understood as follows: “a historically contingent sequence of events is one in which the prior states are necessary or strongly necessary (causal-dependence version), but insufficient (unpredictability version) to bring about the outcome.” Our approach and metrics are compatible with this integrated version, while our results show that deepening the footprint of history can shift the balance between these two aspects of contingency.

### Future directions

Although we quantified the direction and magnitude of phenotypic changes in this work, we have not described them in terms of parallel and convergent evolution. These terms are widely used to describe similar changes, or homoplasy, in two or more independently evolving lineages. Russell Powell notes that many authors attribute homoplasy between closely related lineages to parallel evolution, whereas homoplasy between more distant lineages is attributed to selection’s capacity to drive convergent outcomes [41,42]. Powell suggests using parallelism only when “a developmental homology is the proximate cause of the phenotypic similarity”. He then describes a “screening off” test, adopted from Brandon [43], that could be used to distinguish between parallelism that results from internal, historically contingent constraints and convergence that is driven by external selection. The fine-grained data available with digital organisms should allow us to operationalize that test, or some variant of it, to determine the relative contributions of parallelism and convergence to the homoplastic evolution of specific logic functions in our experiments. A future analysis could use our data to examine how the balance between parallelism and convergence changes with the depth of the footprint of history.

In this work, we also focused on evolutionary outcomes rather than the specific population-genetic processes that led to those outcomes. To examine such processes, Ostrowski et al. [44] used a two-phase experimental design to measure the contributions of antagonistic pleiotropy and mutation accumulation to niche specialization in digital organisms. Antagonistic pleiotropy occurs when a mutation that is beneficial overall nonetheless impairs some other function or functions; thus, it is fundamentally an adaptive process, albeit with tradeoffs. By contrast, mutation accumulation is a stochastic process that occurs when mutations that are not beneficial nonetheless increase by random drift or genetic linkage (e.g., hitchhiking with a beneficial mutation), leading to the reduction or loss of other functions. In the first phase of their experiment, Ostrowski et al. evolved generalists that used many different resources (performed many logic functions), which served as ancestors for populations that evolved in a single-resource environment during the second phase. They found that most mutations that caused the loss of previously rewarded functions were neutral or detrimental in the new environment, indicating the predominance of mutation accumulation. However, antagonistic pleiotropy also played an important role, as evidenced by an over-representation of mutations along the line of descent that were beneficial in the new environment but caused the loss of previously rewarded functions. These findings showed that both random and adaptive processes played important roles in ecological specialization, while also demonstrating that the contributions of those processes can be measured in Avida. Our experiment did not favor specialization, but instead involved a switch from one generalist environment to one of two other generalist environments: one that increased the number of rewarded functions (Overlapping environment), and the other that rewarded a completely different set of functions (Orthogonal environment). Future analyses of our experiments might investigate the contributions of neutral and adaptive processes to the losses and gains of functions, to understand how their relative importance depends on the depth of history as well as on the selective environment. In the Orthogonal environment, we might expect antagonistic pleiotropy to become more important as the footprint of history deepens, because populations with a deep footprint are performing more unrewarded functions, so that more of their genome is a target for beneficial mutations that cause the loss of previously rewarded functions. In the Overlapping environment, by contrast, a shallow history means that more functions that are rewarded in both the old and new habitats remain available for discovery in Phase II, potentially generating synergistic pleiotropy. We expect synergistic pleiotropy would become less common as history deepens in the Overlapping environment, and it should be rare in the Orthogonal environment regardless of the depth of history.

Our study system has many advantages, as well as some potential limitations when making comparisons with biological systems. Avida was designed specifically to study evolution using digital organisms, and it provides many tools that enabled us to observe the process in great detail. To that end, we have constructed and analyzed a self-contained evolving system, comparable in some ways to the *E. coli* Long-Term Evolution Experiment [17,45–54] and other microbial evolution experiments [3,55–61]. While it is difficult to compare evolving microbes to evolving computer programs, it is useful to recognize features of the digital world that are similar to the biological world, as well as others that are different. Some of the similarities include asexual reproduction (as in most but not all evolution experiments with microbes), the experimental tractability afforded by rapid generations, and the ability to store samples for later analysis. On the other hand, the number of evolving populations in Avida is larger than is feasible in most microbial experiments, whereas the number of individuals per population is generally much larger when using microbes. Microbes also have much larger genomes, while per-site mutation rates are usually set much higher in Avida to offset the effects of smaller populations and genomes. The physiology and ecology of microorganisms and Avidians also undoubtedly differ profoundly. For these reasons, experiments with Avida are not meant to simulate biological evolution in any detail, but rather they offer a powerful way of testing general concepts and exploring questions in evolutionary science writ large. In any case, we hope others might use our experimental design—modified as needed for practicality—to study the effects of varying the depth of history’s footprint on evolution in biological systems.

### Concluding thoughts

In closing, we instantiated and extended Gould’s thought-experiment [1] by “replaying life’s tape” from multiple points in the history of these replicate virtual worlds. More than 40 years after Gould and Lewontin’s famous critique of the “adaptationist programme” [62,63], our results also illustrate some of the complex ways in which adaptation, chance, and history together contribute to the evolutionary process. Last but not least, we think that our experimental design can be employed to study the effects of the depth of history’s footprint in biological as well as digital systems.

## Materials and Methods

### Avida system and experiments

We used Avida (version 2.14) to perform the experiments and output the data. Avida is an open-source platform for experiments with digital organisms; it is freely available at https://github.com/devosoft/avida. Charles Ofria led the development of Avida, and it is maintained by the Digital Evolution Lab at Michigan State University.

We started the Phase I populations with a proto-ancestral clone with a genome length of 100 instructions. It was unable to perform any logic functions. Each genome in Avida is a circular series of computational instructions, with 26 possible instructions at each position. While executing its genomic program, a digital organism can copy its genome; if it performs one or more rewarded functions, it obtains additional resources that allow it to run its program at a faster rate. While copying a genome, mutations occur at random; these include both point mutations, which change one instruction to another, and indels, which insert or delete a single instruction. In both Phases I and II, we set the point-mutation rate to 0.007 per instruction copied and the rate of indels to 0.00005 per instruction copied, with indels also occurring at a rate of 0.1 per post-replication division. In both phases, we set the maximum population size to 3600. When a digital organism completed its replication, the new offspring was placed in a randomly chosen position in the population, and the organism that was previously at that position was eliminated. These random deaths thus introduce genetic drift, which along with mutations generate stochasticity in the evolutionary process. Organisms also died if they executed a total number of instructions that exceeded 20 times their genome length; such deaths typically occur if a mutation caused an organism’s genomic program to cycle indefinitely without reproducing itself.

In Avida, an organism’s fitness is calculated as its rate of energy acquisition divided by the cost of producing an offspring, where that cost is equal to the number of instructions it must execute to copy its genome and divide. Note that an organism’s execution length is distinct from (and greater than) its genome length. The execution length represents the total number of instructions performed by a digital organism prior to reproduction, which involves some instructions being executed many times. Execution of an organism’s genomic program includes not only copying the genome but also performing Boolean logic operations that may increase the organism’s rate of energy acquisition.

As in biology, many mutations are deleterious. A mutation might, for example, damage or destroy the copy loop that allows a digital organism to reproduce; or a mutation might disrupt the performance of a logic function that previously evolved, thus reducing the rate at which that organism acquires the energy that fuels the execution of its genomic program. Many other mutations are neutral, having no effect on an organism’s phenotype. Yet other mutations can be beneficial in various ways. Some mutations may reduce the number of instructions needed to copy the genome, thereby making replication a bit more efficient. Others may enable the performance of one of the logic functions rewarded with additional energy. No single instruction is sufficient to perform any of the rewarded functions. However, a single mutation of just the right type may produce a new function in the right genetic background.

The user-defined environment determines whether performance of a particular function is rewarded; this situation is analogous to the choice of which resources are provided in an evolution experiment with bacteria or yeast. In Avida, the environments are defined by two- and three-input Boolean logic functions. If an organism performs a rewarded function while executing its genomic program, then it will receive a reward in the form of additional energy (in the form of CPU cycles) to execute those instructions. All else equal, this additional energy will allow it to reproduce faster, thereby providing an evolutionary advantage. An organism is rewarded for a given function if it outputs a 32-bit string that precisely matches the string that would be expected if it performed the associated bit-wise logic function on specific input strings.

The three environments used in our study reward different subsets of 9 two-input and 68 three-input Boolean logic functions. We excluded the most difficult two-input function (known as “equals” or EQU), leaving a total of 76 other functions. Following Wagenaar and Adami [33], we ordered this set of 76 functions and split it into two halves, called “even” and “odd”, in which adjacent pairs of functions tend to be of similar difficulty given the genomic instructions available in Avida [33]. The Phase I environment rewarded the 38 even functions. In Phase II, the “Overlapping” environment rewarded the same 38 even functions along with the 38 odd functions. The Orthogonal environment, by contrast, rewarded only the 38 odd functions, which were not rewarded during Phase I. The rewards for an organism that can perform two or more logic functions are multiplicative. The specific multipliers for each function, along with other experimental settings, are provided in the configuration and README files at the github repository for this paper: https://https://github.com/bundyjay/Avida_footprint_of_history.

### Analysis

In nested experimental designs like the one that we employed, the contributions of chance and history to the evolution of a trait are estimated using variance components [3,33]. The contribution of chance reflects the variance among Phase II replicate populations derived from the same Phase I progenitor, whereas the contribution of history reflects the additional variation (above that due to chance) associated with the different Phase I progenitors. Both chance and history reflect random, as opposed to fixed, sources of variation. We then constructed 95% confidence intervals around the point estimates using the chi-square distribution [64]. We transformed the variance estimates and their confidence limits by taking the square root of the values, so that the scaling is directly comparable to adaptation [3,33]. We estimated the contribution of adaptation as the mean difference in trait values between each set of 100 Phase II populations and their corresponding progenitors [3,33]. We constructed 95% confidence intervals using the *t* distribution with 99 degrees of freedom (based on the number of Phase II populations). We measured the contributions of adaptation, chance, and history separately for the sets of populations founded from progenitors sampled at three points during Phase I (20,000, 100,000, and 500,000 updates), along with their confidence limits. This approach allowed us to examine how the footprint of history from Phase I affected evolution in both Phase II environments (Figs. 7 & 8). We also measured the contributions of adaptation, chance, and history throughout Phase II, and we show these complete trajectories without their associated confidence limits for visual clarity (Fig. 9 and S8 Fig.).

Owing to the several order-of-magnitude changes in fitness over time, we log_10_-transformed fitness values prior to all analyses. For consistency, we similarly transformed genome lengths, although they varied much less. We performed all data analyses using Python (version 3).

## Acknowledgments

We dedicate this paper to the memory of George Gilchrist, who was the NSF program officer for the BEACON Center for the Study of Evolution in Action. George’s dedication, scholarship, and advice are greatly missed. We thank Louise Mead and Arend Hintze for valuable discussions, and members of the Ofria and Lenski labs for their collegial input. This research received partial support from the BEACON Center (NSF cooperative agreement DBI-0939454) along with fellowships and the John Hannah endowment from Michigan State University.

## Supporting Information

**S1 Fig.**
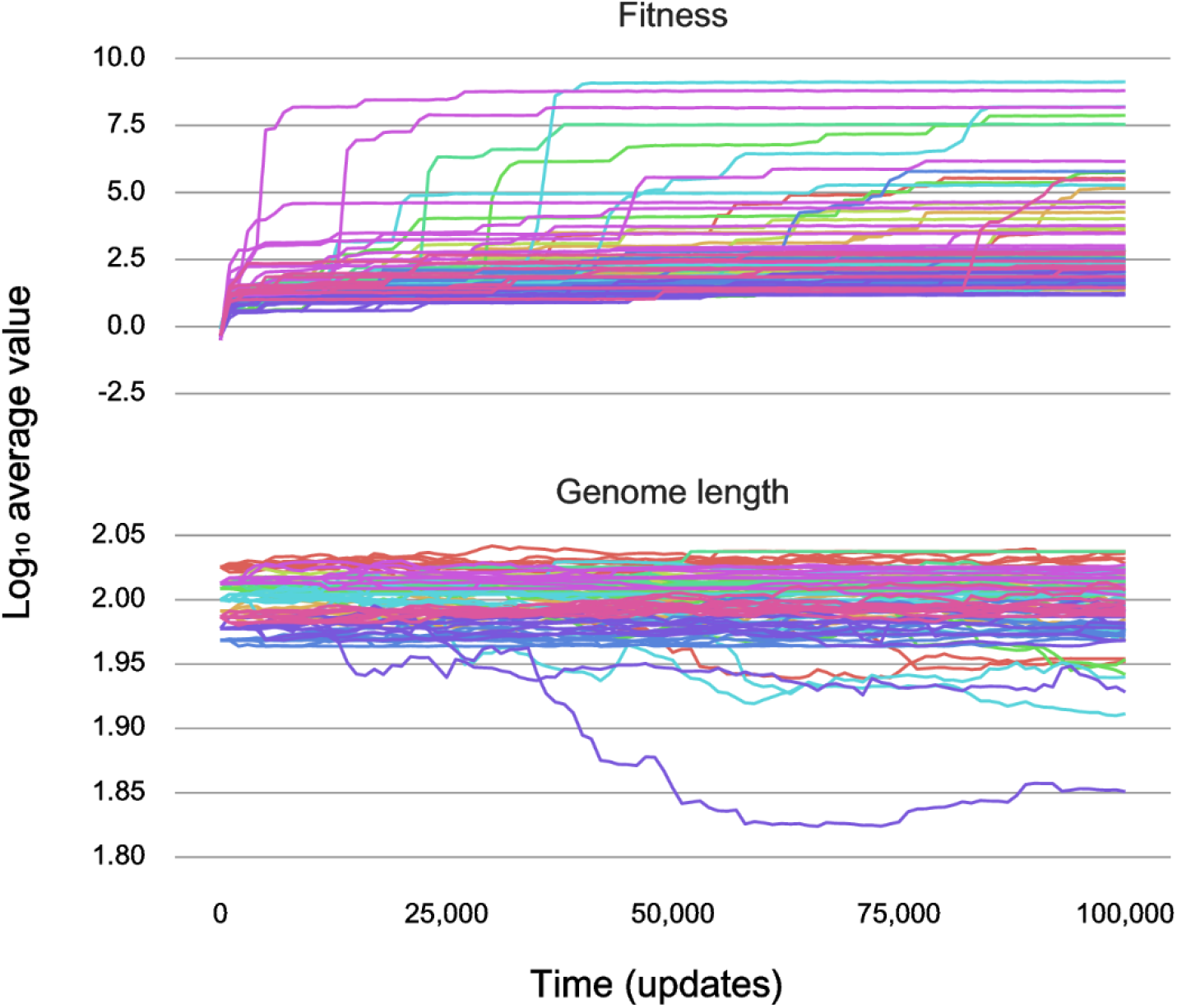
Population averages for fitness and genome length during Phase II in the Orthogonal environment for populations with a shallow history. Average fitness (top) and genome length (bottom) measured every 1,000 updates for 10 replicate populations derived from a single founder from each Phase I population, as they evolved for 100,000 updates in the new Phase II environment. These populations have a shallow history, as their founders evolved for only 20,000 updates in the ancestral environment. The new environment rewarded performance of a set of 38 logic functions that do not overlap with those rewarded in the ancestral environment.

**S2 Fig.**
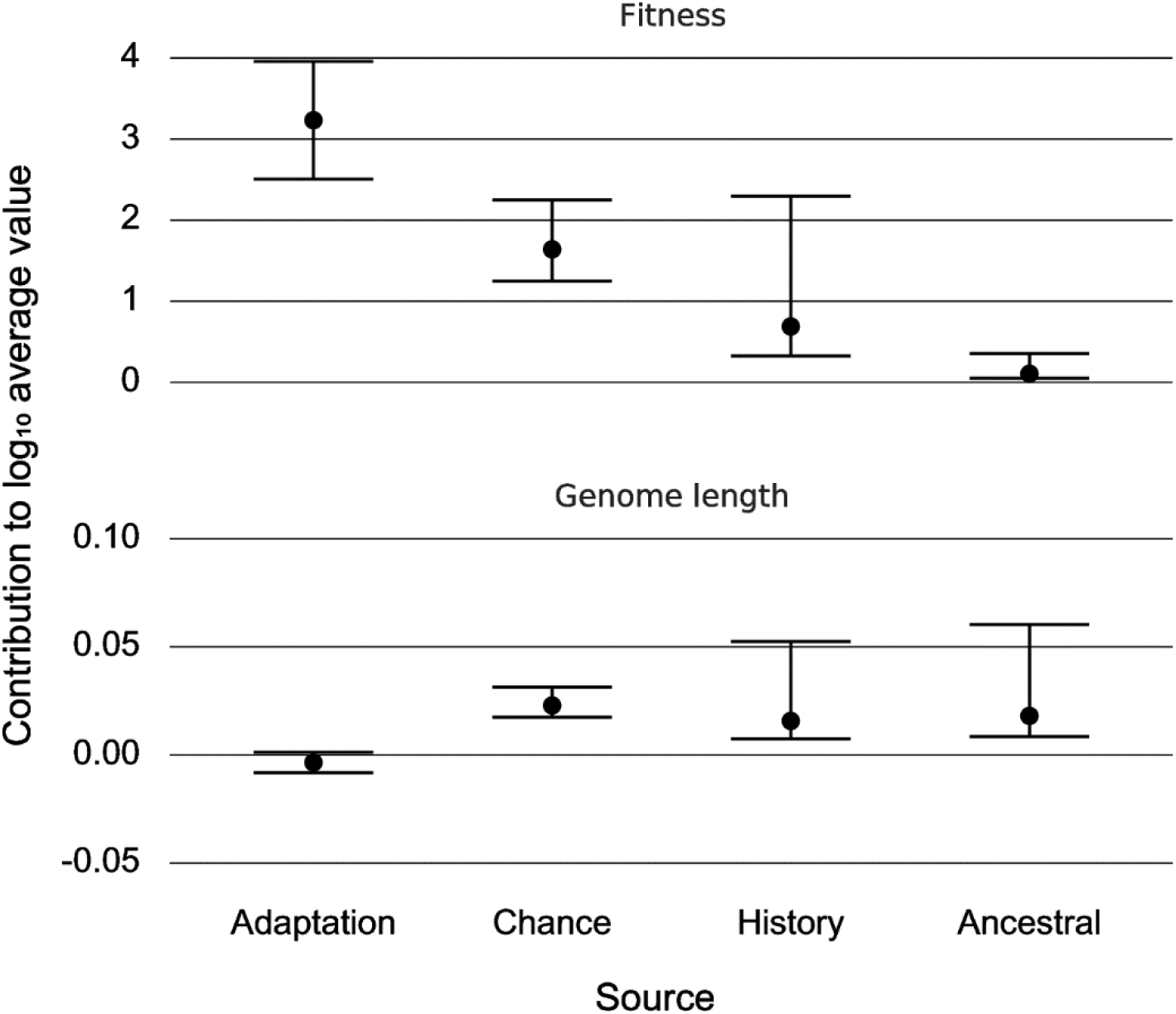
End-of-experiment effects of adaptation, chance, and history in the Orthogonal environment with shallow history. Contributions of adaptation, chance, and history to the evolution of fitness (top) and genome length (bottom) based on end-of-experiment average values for populations with a shallow history in Phase I that evolved in the Orthogonal environment during Phase II. Error bars show 95% confidence limits for the associated contributions, along with the ancestral variation at the start of Phase II.

**S3 Fig.**
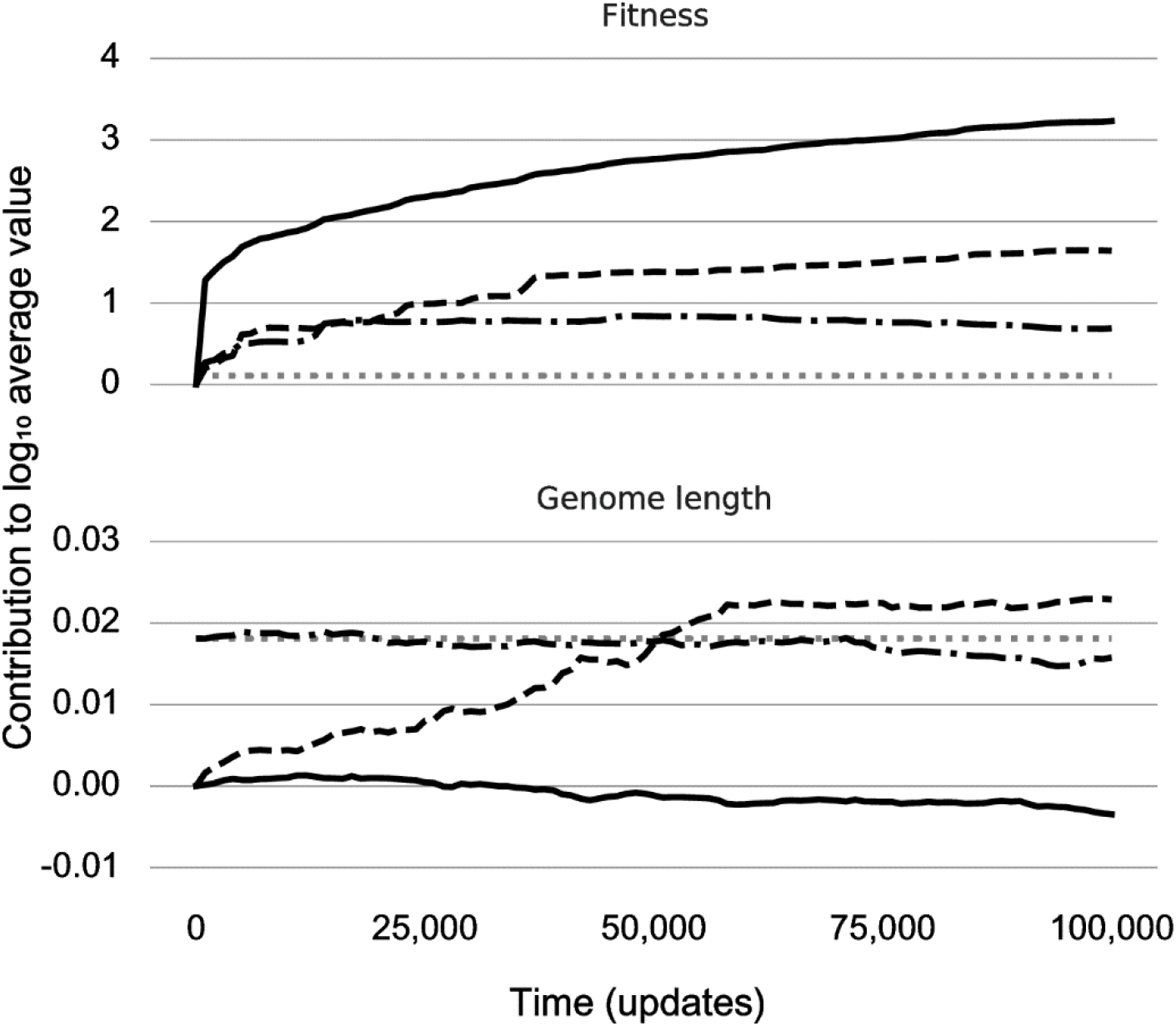
Trajectories of adaptation, chance, and history in the Orthogonal environment with shallow history. Contributions of adaptation (solid line), chance (dashed line), and history (dash-dotted line) to the evolution of fitness (top) and genome length (bottom) measured every 1,000 updates during Phase II. This plot shows the ancestral variance (dotted line) for comparison.

**S4 Fig.**
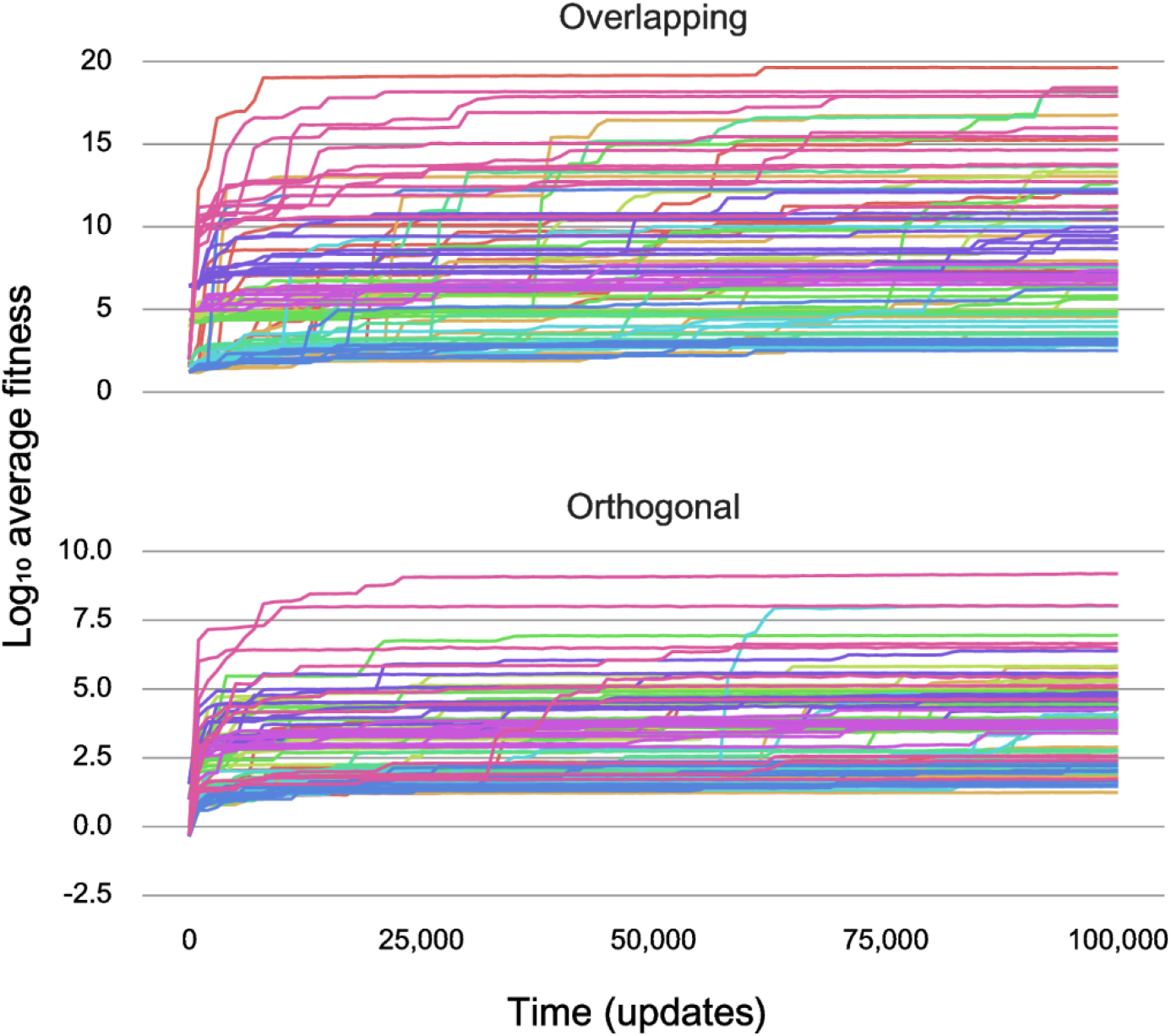
Population averages for fitness during Phase II with intermediate history. Average fitness in the Overlapping (top) and Orthogonal (bottom) environments measured every 1,000 updates for 10 replicate populations derived from a single founder from each Phase I population, as they evolved for 100,000 updates in the new Phase II environment. These populations have an intermediate history, as their founders evolved for 100,000 updates in the ancestral environment.

**S5 Fig.**
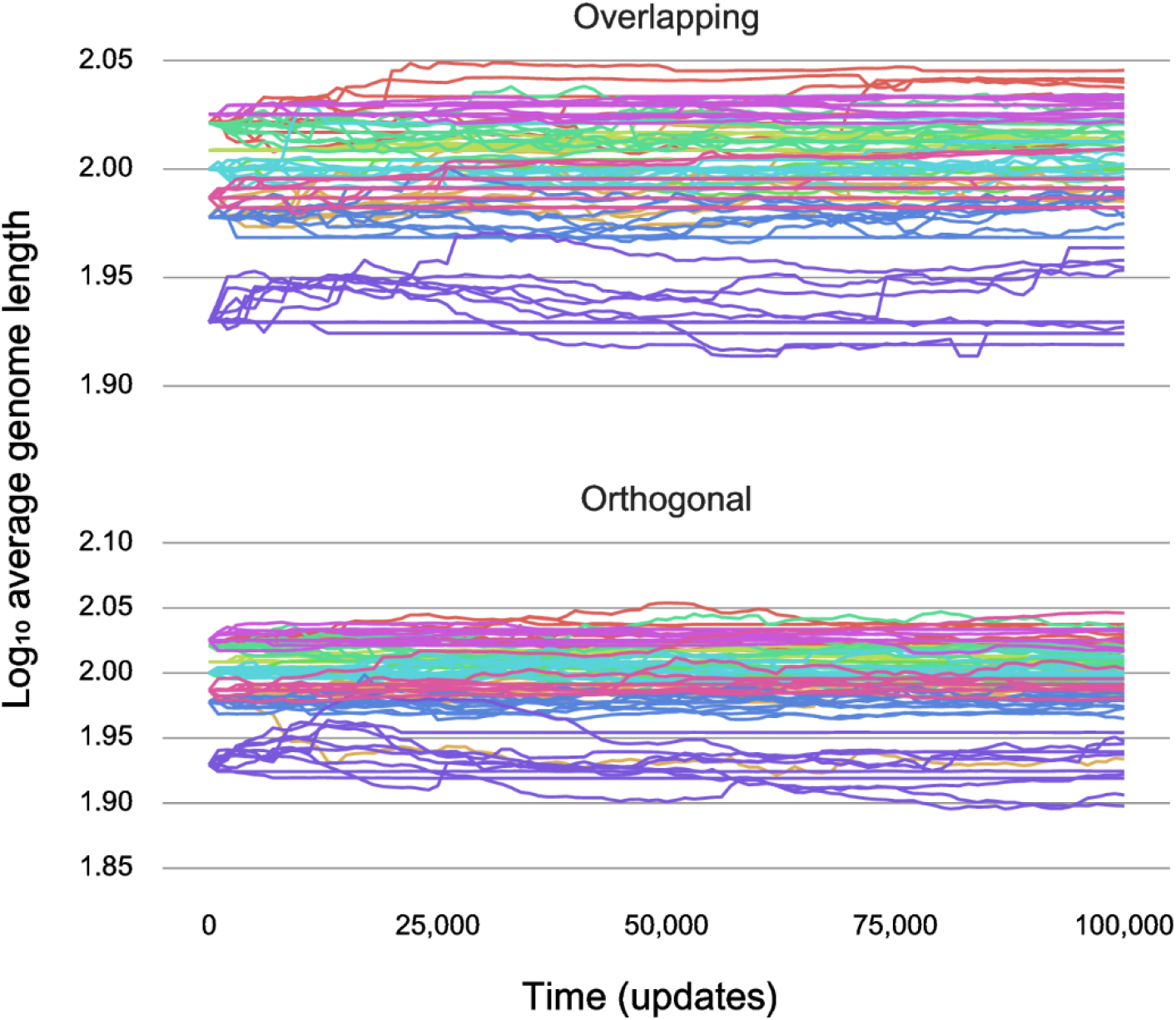
Population averages for genome length during Phase II with intermediate history. Average genome length in the Overlapping (top) and Orthogonal (bottom) environments measured every 1,000 updates for 10 replicate populations derived from a single founder from each Phase I population, as they evolved for 100,000 updates in the new Phase II environment. These populations have an intermediate history, as their founders evolved for 100,000 updates in the ancestral environment.

**S6 Fig.**
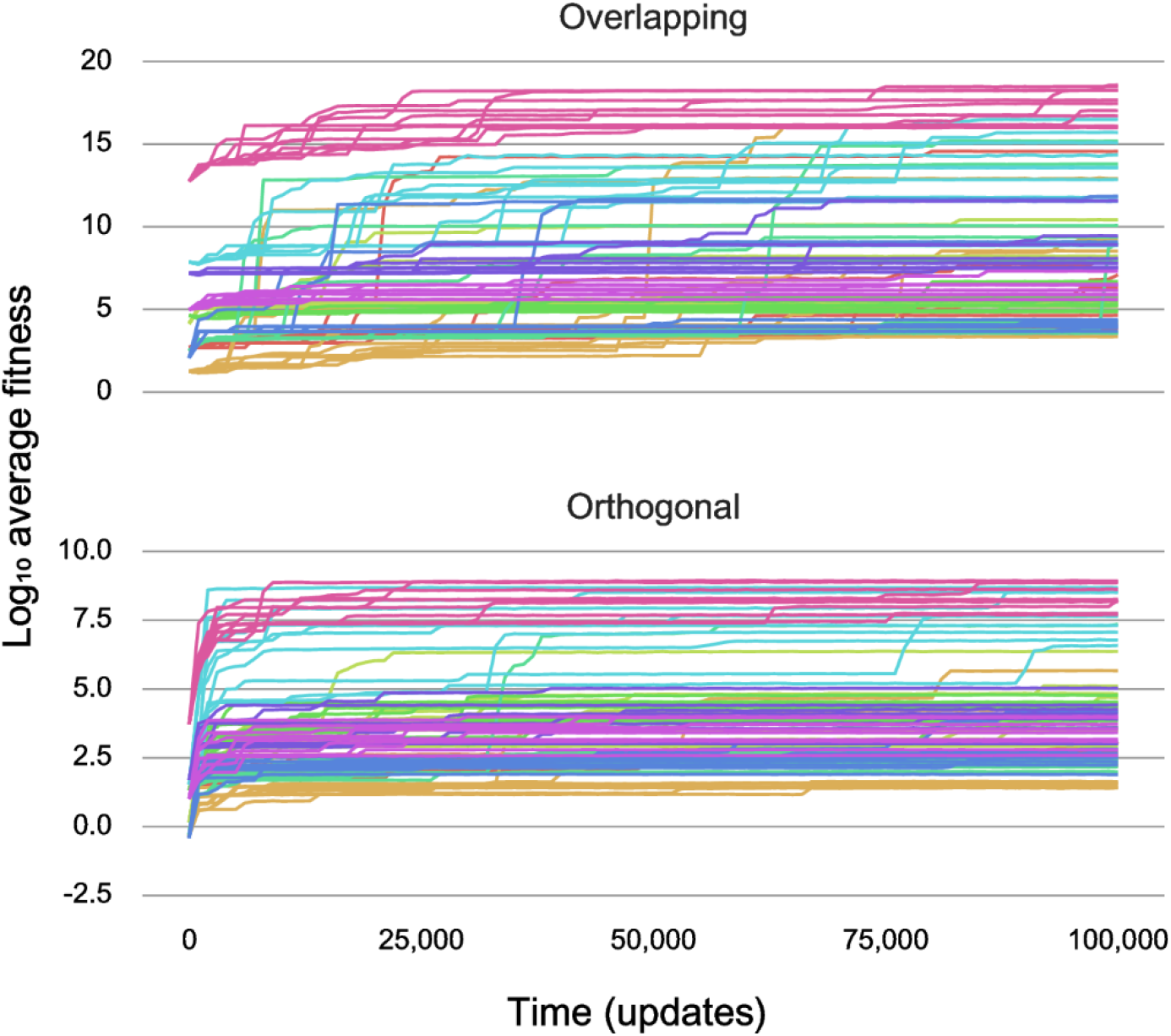
Population averages for fitness during Phase II with deep history. Average fitness in the Overlapping (top) and Orthogonal (bottom) environments measured every 1,000 updates for 10 replicate populations derived from a single founder from each Phase I population, as they evolved for 100,000 updates in the new Phase II environment. These populations have a deep history, as their founders evolved for 500,000 updates in the ancestral environment.

**S7 Fig.**
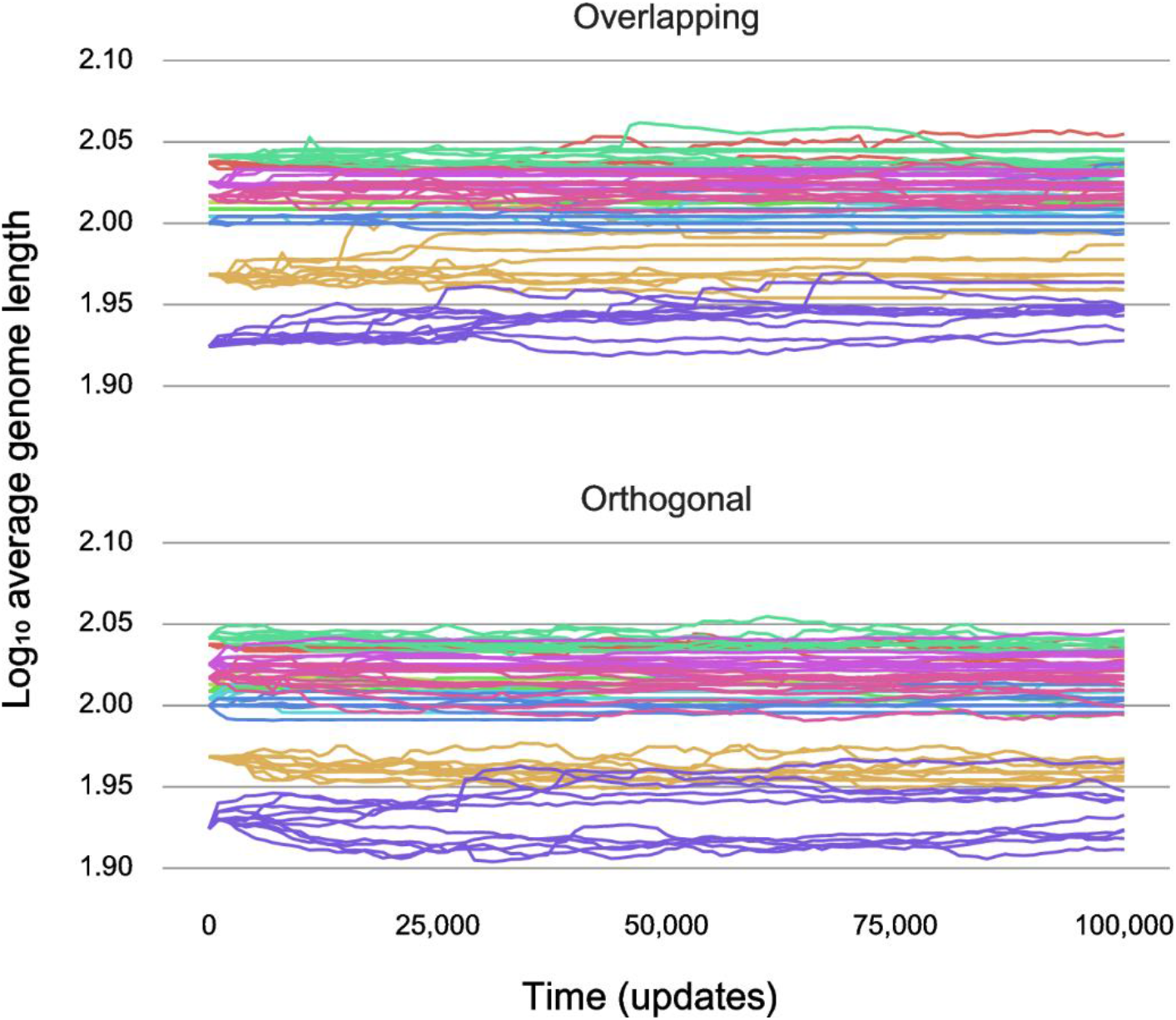
Population averages for genome length during Phase II with deep history. Average genome length in the Overlapping (top) and Orthogonal (bottom) environments measured every 1,000 updates for 10 replicate populations derived from a single founder from each Phase I population, as they evolved for 100,000 updates in the new Phase II environment. These populations have a deep history, as their founders evolved for 500,000 updates in the ancestral environment.

**S8 Fig.**
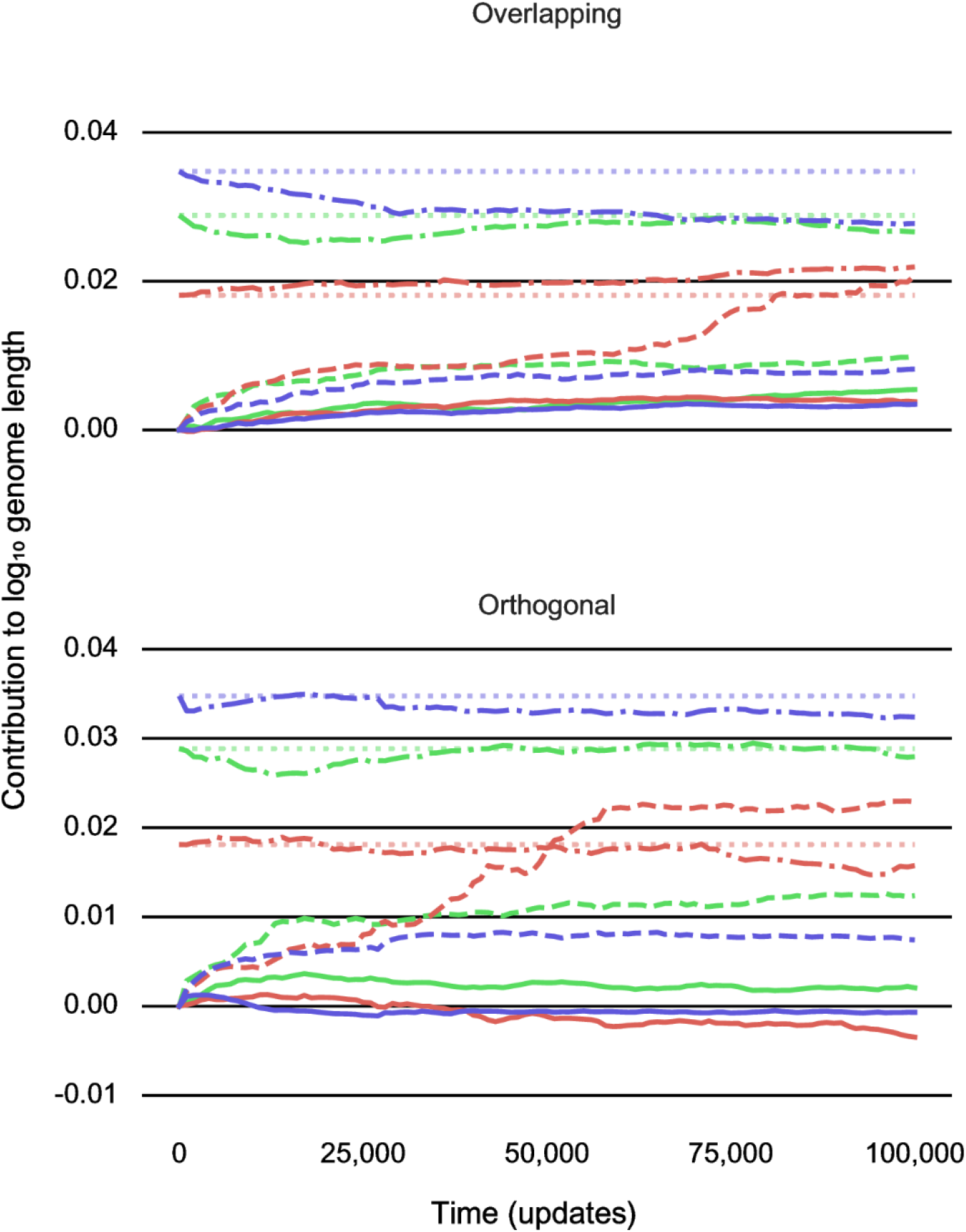
Trajectories for the contributions of adaptation, chance, and history to genome length during Phase II. Contributions of adaptation (solid lines), chance (dashed lines), and history (dash-dotted lines) to the evolution of genome length in the Overlapping (top) and Orthogonal (bottom) environments measured every 1,000 updates during Phase II for populations with shallow (red), intermediate (green), and deep (blue) Phase I histories. This plot shows the ancestral variances (dotted lines) for comparison.

## References

1. Gould SJ. Wonderful Life. W. W. Norton; 1990.

2. Blount ZD, Lenski RE, Losos JB. Contingency and determinism in evolution: replaying life’s tape. Science. 2018; 362(6415):1–10.

3. Travisano M, Mongold JA, Bennett AF, Lenski RE. Experimental tests of the roles of adaptation, chance, and history in evolution. Science. 1995; 267(5194):87–90.

4. Losos JB, Jackman TR, Larson A, de Queiroz K, Rodriguez-Schettino L. Contingency and determinism in replicated adaptive radiations of island lizards. Science. 1998; 279(5359):2115–2118.

5. Teotónio H, Rose MR. Variation in the reversibility of evolution. Nature. 2000; 408(6811):463–466.

6. Emerson SB. A macroevolutionary study of historical contingency in the fanged frogs of Southeast Asia. Biol J Linn Soc Lond. 2001; 73(1):139–151.

7. Vanhooydonck B, Irschick DJ. Is evolution predictable? Evolutionary relationships of divergence in ecology, performance and morphology in Old and New World lizard radiations. Aerts P, D’Août K, Herrel A, Van Damme R, editors. Topics in Functional and Ecological Vertebrate Morphology. 2002; 191–204.

8. Morris SC. Life’s Solution. Cambridge Univ Press; 2003.

9. Joshi A, Castillo RB, Mueller LD. The contribution of ancestry, chance, and past and ongoing selection to adaptive evolution. J Genet. 2003; 82(3):147–162.

10. Ortlund EA, Bridgham JT, Redinbo MR, Thornton JW. Crystal structure of an ancient protein: evolution by conformational epistasis. Science. 2007; 317(5844):1544–1548.

11. Keller SR, Taylor DR. History, chance and adaptation during biological invasion: separating stochastic phenotypic evolution from response to selection. Ecol Lett. 2008; 11(8):852–866.

12. Dick MH, Lidgard S, Gordon DP, Mawatari SF. The origin of *Ascophoran bryozoans* was historically contingent but likely. Proc R Soc B. 2009; 276(1670):3141–3148.

13. Conway Morris S. Evolution: like any other science it is predictable. Phil Trans R Soc B. 2010; 365(1537):133–145.

14. Meyer JR, Dobias DT, Weitz JS, Barrick JE, Quick RT, Lenski RE. Repeatability and contingency in the evolution of a key innovation in phage lambda. Science. 2012; 335(6067):428–432.

15. Harms MJ, Thornton JW. Historical contingency and its biophysical basis in glucocorticoid receptor evolution. Nature. 2014; 512(7513):203–207.

16. Starr TN, Picton LK, Thornton JW. Alternative evolutionary histories in the sequence space of an ancient protein. Nature. 2017; 549(7672):409–413.

17. Lenski RE, Rose MR, Simpson SC, Tadler SC. Long-term experimental evolution in *Escherichia coli*. I. Adaptation and divergence during 2,000 generations. Am Nat. 1991; 138(6):1315–1341.

18. Flores-Moya A, Costas E, López-Rodas V. Roles of adaptation, chance and history in the evolution of the dinoflagellate *Prorocentrum triestinum*. Naturwissenschaften. 2008; 95(8):697–703.

19. Flores-Moya A, Rouco M, García-Sánchez MJ, García-Balboa C, González R, Costas E, et al. Effects of adaptation, chance, and history on the evolution of the toxic dinoflagellate *Alexandrium minutum* under selection of increased temperature and acidification. Ecol Evol. 2012; 2(6):1251–1259.

20. Pérez-Zaballos FJ, Ortega-Mora LM, Álvarez-García G, Collantes-Fernández E, Navarro-Lozano V, García-Villada L, et al. Adaptation of *Neospora caninum* isolates to cell-culture changes: an argument in favor of its clonal population structure. J Parasitol. 2005; 507–510.

21. Rouco M, López-Rodas V, Flores-Moya A, Costas E. Evolutionary changes in growth rate and toxin production in the cyanobacterium *Microcystis aeruginosa* under a scenario of eutrophication and temperature increase. Microb Ecol. 2011; 62(2):265–273.

22. Rebolleda‐Gómez M, Travisano M. Adaptation, chance, and history in experimental evolution reversals to unicellularity. Evolution. 2019; 73(1):73–83.

23. Santos-Lopez A, Marshall CW, Welp AL, Turner C, Rasero J, Cooper VS. The roles of history, chance, and natural selection in the evolution of antibiotic resistance. bioRxiv. 2020; 2020.07.22.216465.

24. Burch CL, Lin C. Evolvability of an RNA virus is determined by its mutational neighbourhood. Nature. 2000; 406(6796):625–628.

25. Teotónio H, Chelo IM, Bradić M, Rose MR, Long AD. Experimental evolution reveals natural selection on standing genetic variation. Nat Genet. 2009; 41(2):251–257.

26. Bedhomme S, Lafforgue G, Elena SF. Genotypic but not phenotypic historical contingency revealed by viral experimental evolution. BMC Evol Biol. 2013; 13(1):46.

27. Kryazhimskiy S, Rice DP, Jerison ER, Desai MM. Global epistasis makes adaptation predictable despite sequence-level stochasticity. Science. 2014; 344(6191):1519–1522.

28. Spor A, Kvitek DJ, Nidelet T, Martin J, Legrand J, Dillmann C, et al. Phenotypic and genotypic convergences are influenced by historical contingency and environment in yeast. Evolution. 2014; 68(3):772–790.

29. Simões P, Fragata I, Seabra SG, Faria GS, Santos MA, Rose MR, et al. Predictable phenotypic, but not karyotypic, evolution of populations with contrasting initial history. Sci Reports. 2017; 7(1):913.

30. Taylor T, Hallam J. Replaying the tape: An investigation into the role of contingency in evolution. In: Artificial Life VI: Proceedings of the Sixth International Conference on Artificial Life. MIT Press; 1998. p. 256–265.

31. Yedid G, Bell G. Macroevolution simulated with autonomously replicating computer programs. Nature. 2002; 420(6917):810–812.

32. Lenski RE, Ofria C, Pennock RT, Adami C. The evolutionary origin of complex features. Nature. 2003; 423(6936):139–144.

33. Wagenaar DA, Adami C. Influence of chance, history, and adaptation on digital evolution. Artificial Life. 2004; 10(2):181–190.

34. Braught G, Dean A. The effects of learning on the roles of chance, history and adaptation in evolving neural networks. In: Australian Conference on Artificial Life. Berlin, Heidelberg: Springer; 2007; p. 201–211.

35. Yedid G, Ofria CA, Lenski RE. Historical and contingent factors affect re-evolution of a complex feature lost during mass extinction in communities of digital organisms. J Evol Biol. 2008; 21(5):1335–1357.

36. Adami C. Introduction to Artificial Life. Springer; 1998.

37. Lenski RE, Ofria C, Collier TC, Adami C. Genome complexity, robustness and genetic interactions in digital organisms. Nature. 1999; 400(6745):661–664.

38. Wilke CO, Wang JL, Ofria C, Lenski RE, Adami C. Evolution of digital organisms at high mutation rates leads to survival of the flattest. Nature. 2001; 412(6844):331–333.

39. Dawkins R. The Blind Watchmaker. W. W. Norton; 2015.

40. Beatty J. Replaying life’s tape. J Philosophy. 2006; 103(7):336–362.

41. Powell R. Is convergence more than an analogy? Homoplasy and its implications for macroevolutionary predictability. Biol Philos. 2007; 22(4):565–578.

42. Powell R. Contingency and convergence in macroevolution: a reply to John Beatty. J Philosophy. 2009; 106(7):390–403.

43. Brandon RN. Adaptation and Environment. Princeton Univ Press; 1990.

44. Ostrowski EA, Ofria C, Lenski RE. Ecological specialization and adaptive decay in digital organisms. Am Nat. 2007; 169(1):E1–20.

45. Lenski RE, Travisano M. Dynamics of adaptation and diversification: a 10,000-generation experiment with bacterial populations. Proc Natl Acad Sci USA. 1994; 91(15):6808–6814.

46. Sniegowski PD, Gerrish PJ, Lenski RE. Evolution of high mutation rates in experimental populations of *E. coli*. Nature. 1997; 387(6634):703–705.

47. Cooper VS, Lenski RE. The population genetics of ecological specialization in evolving *Escherichia coli* populations. Nature. 2000; 407(6805):736–739.

48. Blount ZD, Borland CZ, Lenski RE. Historical contingency and the evolution of a key innovation in an experimental population of *Escherichia coli*. Proc Natl Acad Sci USA. 2008; 105(23):7899–7906.

49. Khan AI, Dinh DM, Schneider D, Lenski RE, Cooper TF. Negative epistasis between beneficial mutations in an evolving bacterial population. Science. 2011; 332(6034):1193–1196.

50. Blount ZD, Barrick JE, Davidson CJ, Lenski RE. Genomic analysis of a key innovation in an experimental *Escherichia coli* population. Nature. 2012; 489(7417):513–518.

51. Good BH, McDonald MJ, Barrick JE, Lenski RE, Desai MM. The dynamics of molecular evolution over 60,000 generations. Nature. 2017; 551(7678):45–50.

52. Card KJ, LaBar T, Gomez JB, Lenski RE. Historical contingency in the evolution of antibiotic resistance after decades of relaxed selection. PLoS Biol. 2019; 17(10).

53. Blount ZD, Maddamsetti R, Grant NA, Ahmed ST, Jagdish T, Baxter JA, et al. Genomic and phenotypic evolution of *Escherichia coli* in a novel citrate-only resource environment. Elife. 2020; 9:e55414.

54. Grant NA, Maddamsetti R, Lenski RE. Maintenance of metabolic plasticity despite relaxed selection in a long-term evolution experiment with *Escherichia coli*. Am Nat. 2021; in press.

55. Rainey PB, Travisano M. Adaptive radiation in a heterogeneous environment. Nature. 1998; 394(6688):69–72.

56. Kawecki TJ, Lenski RE, Ebert D, Hollis B, Olivieri I, Whitlock MC. Experimental evolution. Trends Ecol Evol. 2012; 27(10):547–560.

57. Ratcliff WC, Denison RF, Borrello M, Travisano M. Experimental evolution of multicellularity. Proc Natl Acad Sci USA. 2012; 109(5):1595–1600.

58. Barrick JE, Lenski RE. Genome dynamics during experimental evolution. Nat Rev Gen. 2013; 14(12):827–839.

59. Lang GI, Rice DP, Hickman MJ, Sodergren E, Weinstock GM, Botstein D, et al. Pervasive genetic hitchhiking and clonal interference in forty evolving yeast populations. Nature. 2013; 500(7464):571–574.

60. Levy SF, Blundell JR, Venkataram S, Petrov DA, Fisher DS, Sherlock G. Quantitative evolutionary dynamics using high-resolution lineage tracking. Nature. 2015; 519(7542):181–186.

61. Johnson MS, Gopalakrishnan S, Goyal J, Dillingham ME, Bakerlee CW, Humphrey PT, et al. Phenotypic and molecular evolution across 10,000 generations in laboratory budding yeast populations. Elife. 2021; 10:e63910.

62. Gould SJ, Lewontin RC. The spandrels of San Marco and the Panglossian paradigm: a critique of the adaptationist programme. Proc R Soc B. 1979; 205(1161):581–598.

63. Price RM, Perez KE. Beyond the adaptationist legacy: updating our teaching to include a diversity of evolutionary mechanisms. Am Biol Teacher. 2016; 78(2):101–108.

64. Sokal RR, Rohlf J. Biometry. 3rd ed. New York, NY: W. H. Freeman; 1994.

